# A developmental resource mapping tubulin isotype deployment during corneal morphogenesis

**DOI:** 10.64898/2026.02.19.706651

**Authors:** R Ramarapu, WR Stoehr, M Miesen, S. Border, SM Thomasy, CD Rogers

**Author notes:** These authors contributed to this manuscript equally (R Ramarapu, WR Stoehr).

## Abstract

Microtubules are core cytoskeletal polymers that support epithelial organization, polarity, and tissue morphogenesis. They are composed of α– and β-tubulin heterodimers encoded by multigene families that generate closely related but distinct isotypes. Although several tubulin isotypes have been implicated in ocular development and disease, their spatial organization during corneal morphogenesis remains poorly defined. Here we establish a resource visualizing and quantifying tubulin isotype localization during corneal development using the chick embryo. Comparative amino acid sequence analysis demonstrates that chick and human α– and β-tubulin isotypes are highly conserved within structural and catalytic domains, with divergence concentrated in C-terminal regions normally associated with post-translational modification sites. Relating sequence features to tissue organization, we performed longitudinal immunohistochemical analysis of five tubulin isotypes across corneal development. These analyses reveal distinct and dynamic patterns of isotype enrichment along apicobasal axes, as well as stage-dependent redistribution during epithelial maturation and endothelial differentiation. TUBA5/TUBA4A displays particularly restricted localization, including enrichment in migratory corneal stromal progenitors and the developing corneal epithelium and endothelium. Together, these data provide a conserved spatial framework for tubulin isotype organization during corneal morphogenesis and establish the chick embryo as a tractable model for investigating isotype-specific microtubule organization during epithelial development.

**SIGNIFICANCE STATEMENT:** This study provides a spatial and developmental resource that visualizes when and where distinct tubulin isotypes are deployed during corneal morphogenesis.

## INTRODUCTION

The cornea is the transparent anterior tissue of the eye and plays a central role in light refraction, optical clarity, and protection of the ocular surface. Its development requires the coordinated morphogenesis of three distinct layers, the stratified epithelium, the collagen-rich stroma, and the corneal endothelial monolayer, each of which emerges through tightly regulated processes of cell migration, polarization, differentiation, and tissue-scale organization (Kivela and Uusitalo 1998, Eghrari, Riazuddin et al. 2015, Chen and Dong 2022, Ramarapu, Stoehr et al. 2026). Underlying these events is a dynamic cytoskeletal network that integrates mechanical support with intracellular transport and signaling (Weinreb and Ryder 1990, Su, Desikan et al. 2025). Establishing a resource would enable comparisons across tissues and species and provide a framework for investigating how microtubule diversity contributes to epithelial organization and morphogenesis.

Microtubules are a core component of this cytoskeletal network and are essential for epithelial organization, directional migration, vesicle trafficking, and force generation (Laan, Husson et al. 2008, Garcin and Straube 2019). They are assembled from α– and β-tubulin heterodimers, two highly conserved protein families encoded by multiple genes that give rise to distinct isotypes. Although tubulin isotypes differ by only a small number of amino acids, accumulating evidence indicates that these differences, together with extensive post-translational modifications (PTMs), generate functionally specialized microtubule populations across tissues and developmental contexts (Hausrat, Radwitz et al. 2021, Moutin, Bosc et al. 2021, Bar, Popp et al. 2022, Carmona, Marinho et al. 2023). This concept, often referred to as the “tubulin code,” proposes that isotype composition and PTMs tune microtubule stability, interactions with motors and microtubule-associated proteins (MAPs), and intracellular organization. Importantly, these two sources of microtubule diversity are not necessarily independent. Many PTMs occur within the highly divergent C-terminal tails of α– and β-tubulins, such that isotype-specific sequence differences can alter the availability of modification sites, recognition by PTM enzymes, and interactions with microtubule motors and MAPs (Janke and Magiera 2020, Lu and Zheng 2022, Bao, Dorig et al. 2023, Fu, Yan et al. 2023). Thus, developmental regulation of tubulin isotype expression has the potential to alter both the molecular composition of the microtubule lattice and the repertoire of PTMs and interacting proteins available to that lattice.

Spatiotemporal expression of tubulin isotypes has been most extensively characterized in the nervous system, where mutations in both α– and β-tubulin genes cause a spectrum of developmental disorders collectively termed tubulinopathies. These conditions routinely affect the central nervous system and include cortical malformations, axon guidance defects, epilepsy, hypomyelination, and corpus callosum abnormalities (Aiken, Buscaglia et al. 2017, Schroter, Doring et al. 2021, Schroter, Popp et al. 2022, Ganne, Balasubramaniam et al. 2023, Torella, Ricca et al. 2023, Benkirane, Bonhomme et al. 2024, Wan, Zhou et al. 2024). By contrast, the roles of tubulin isotypes in ocular development, specifically in the cornea, remain limited. Emerging clinical and experimental evidence suggests that this gap in knowledge is noteworthy (Table 1). Mutations in *TUBB3* underlie congenital fibrosis of the extraocular muscles, a disorder characterized by impaired ocular motility and structural abnormalities (Chew, Balasubramanian et al. 2013, Whitman, Andrews et al. 2016). *De novo* mutations in *TUBA3D* have been associated with keratoconus, which affects cornea shape and function (Hao, Chen et al. 2017). Additionally, recent work has linked microtubule dysregulation, including altered *TUBB4A* expression, to Fuchs’ endothelial corneal dystrophy downstream of TCF4 activity (Yan, Mehta et al. 2024). Together, these findings point to previously underappreciated roles for tubulin isotypes in corneal morphogenesis and homeostasis.

**Table 1.**
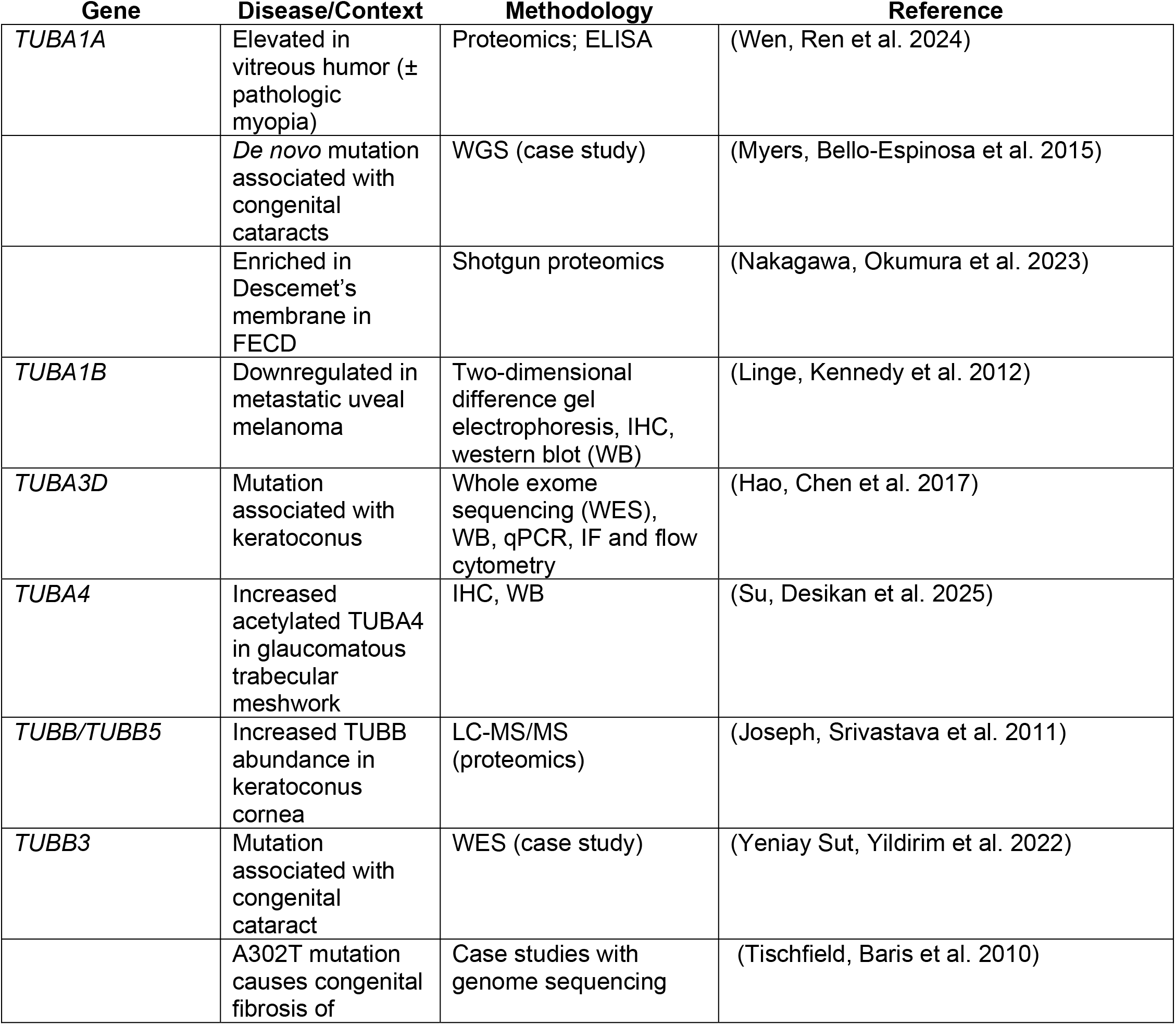
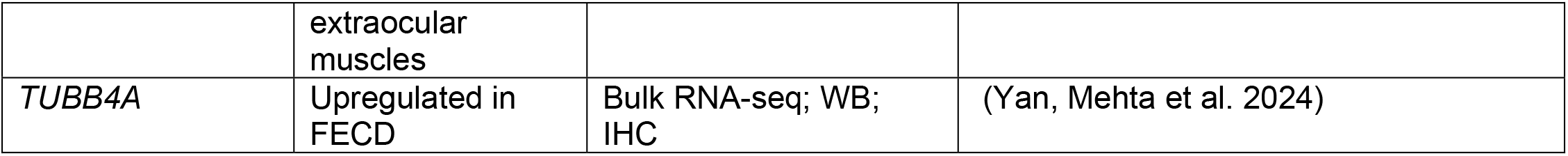
Tubulin isotypes associated with ocular disease.

| Gene | Disease/Context | Methodology | Reference |
| --- | --- | --- | --- |
| *TUBA1A* | Elevated in vitreous humor (± pathologic myopia) | Proteomics; ELISA | (Wen, Ren et al. 2024) |
|  | *De novo* mutation associated with congenital cataracts | WGS (case study) | (Myers, Bello-Espinosa et al. 2015) |
|  | Enriched in Descemet's membrane in FECD | Shotgun proteomics | (Nakagawa, Okumura et al. 2023) |
| *TUBA1B* | Downregulated in metastatic uveal melanoma | Two-dimensional difference gel electrophoresis, IHC, western blot (WB) | (Linge, Kennedy et al. 2012) |
| *TUBA3D* | Mutation associated with keratoconus | Whole exome sequencing (WES), WB, qPCR, IF and flow cytometry | (Hao, Chen et al. 2017) |
| *TUBA4* | Increased acetylated TUBA4 in glaucomatous trabecular meshwork | IHC, WB | (Su, Desikan et al. 2025) |
| *TUBB/TUBB5* | Increased TUBB abundance in keratoconus cornea | LC-MS/MS (proteomics) | (Joseph, Srivastava et al. 2011) |
| *TUBB3* | Mutation associated with congenital cataract | WES (case study) | (Yeniay Sut, Yildirim et al. 2022) |
|  | A302T mutation causes congenital fibrosis of extraocular muscles | Case studies with genome sequencing | (Tischfield, Baris et al. 2010) |
| *TUBB4A* | Upregulated in FECD | Bulk RNA-seq; WB; IHC | (Yan, Mehta et al. 2024) |

The chick embryo provides a powerful system to address the role of tubulin function during development. Chick corneal development is highly accessible, follows a well-defined temporal sequence, and is morphogenetically conserved with human ocular development (Ritchey, Code et al. 2011, Lwigale 2015). Moreover, the chick has long served as a model for studying periocular mesenchyme migration, epithelial-mesenchymal interactions, and corneal layer specification. However, a systematic resource describing tubulin isotype conservation and spatiotemporal localization during corneal development is currently lacking.

Here, we present a combined computational and experimental resource characterizing the spatiotemporal localization of several α– and β-tubulin isotypes in the developing chick cornea. We first perform a comparative sequence analysis of chick and human tubulin isotypes to assess conservation, phylogenetic relationships, and predicted PTM availability. We then provide a longitudinal spatiotemporal map with quantitative signal analysis of five tubulin isotypes (TUBA1A, TUBA1B, TUBA5/TUBA4A, TUBB1/TUBB2A, and TUBB4/TUBB3) using immunohistochemistry (IHC) across key stages of corneal development. Together, these data establish the chick as a robust model for studying isotype-specific microtubule biology in the cornea as well as other epithelial and mesenchymal tissues and provide a foundational framework for future functional and disease-oriented studies.

## RESULTS

### Conservation of tubulin isotype sequences between chick and human

To evaluate the extent to which chick tubulin protein sequences are conserved with their human counterparts, and to identify sequence features that may underlie isotype-specific functional differences, we aligned all RefSeq α– and β-tubulin protein sequences from *Homo sapiens* (human) and *Gallus gallus* (chicken) (Table 2). Analyses were performed separately for α– and β-tubulins and included multiple sequence alignment, per-residue heterogeneity scoring, phylogenetic reconstruction, and pairwise identity comparisons (Fig. 1, 2).

**Figure 1.**
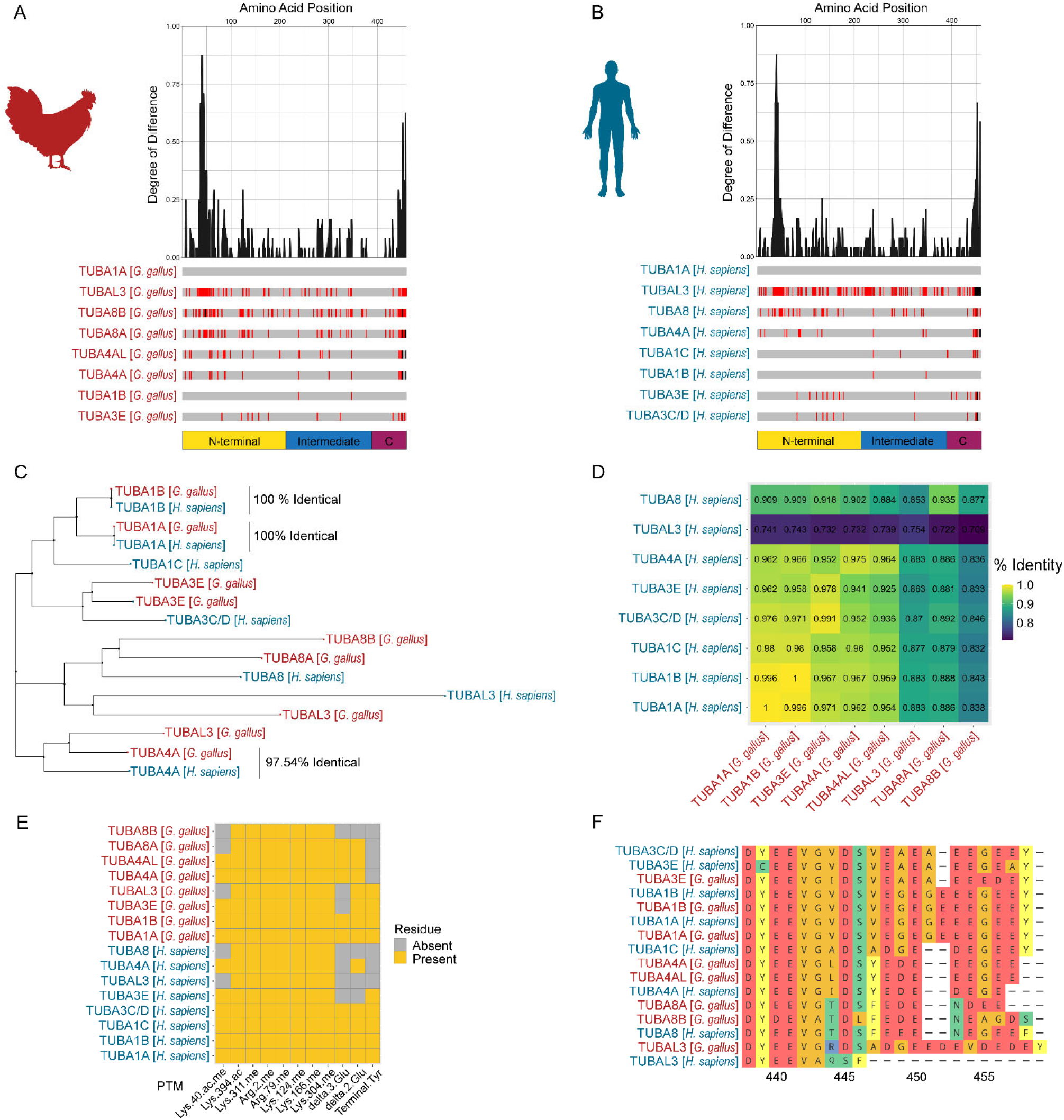
Conservation and divergence of α-tubulin isotypes between chick and human. Chick and human α-tubulin isotypes are highly conserved across core structural domains, with divergence concentrated in C-terminal regions enriched for post-translational modification sites. (A,B) Multiple sequence alignments of α-tubulin isotypes from (A) *Gallus gallus* and (B) *Homo sapiens*. Amino acid mismatches relative to TUBA1A are shown in red, and gaps are shown in black. Plots above each alignment indicate per-residue heterogeneity, calculated as the complement of the conservation score averaged over three-residue bins. (C) Neighbor-joining phylogenetic tree of human and chick α-tubulin isotypes based on amino acid similarity. (D) Pairwise amino acid identity matrix for human and chick α-tubulin isotypes. (E) Distribution of canonical and predicted post-translational modification sites across α-tubulin sequences. Yellow indicates presence of the target residue; grey indicates absence. (F) Alignment of α-tubulin C-terminal tails. Human sequences are shown in blue and chick sequences in red.

**Table 2.**
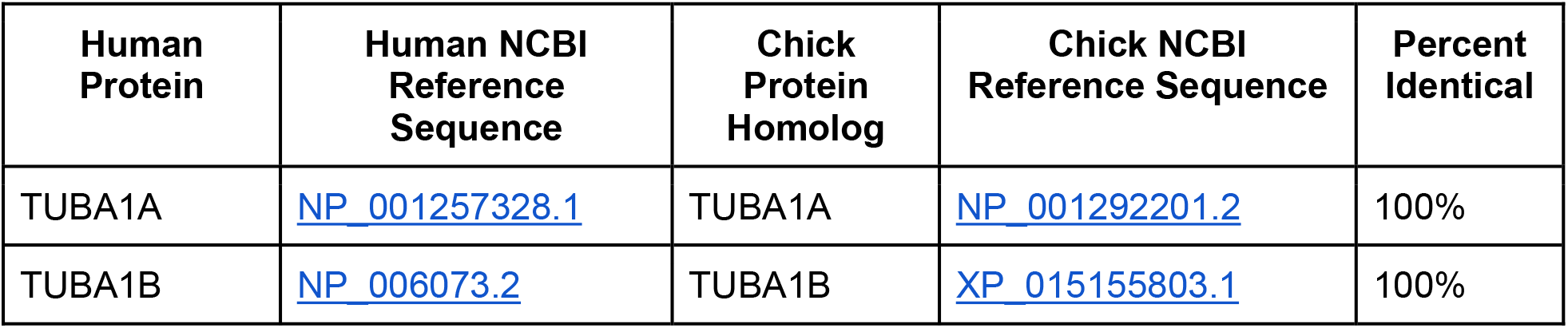
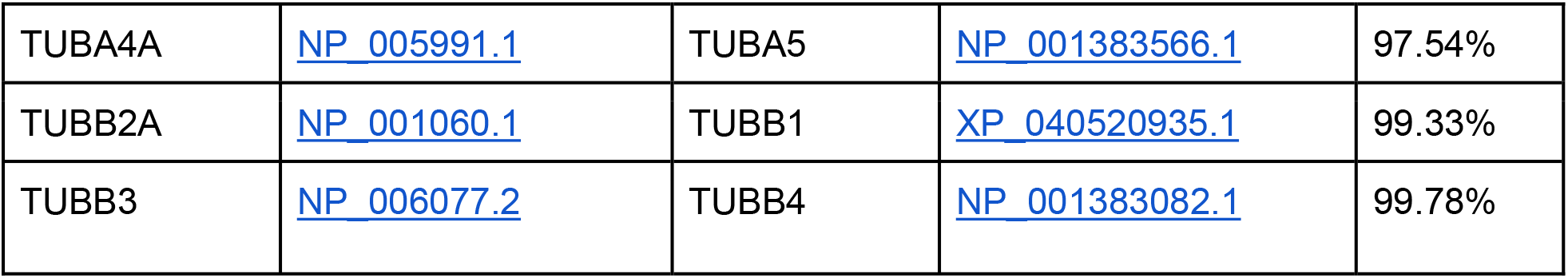
Tubulin Nomenclature.

Multiple sequence alignments revealed a high degree of conservation across both species for all tubulin isotypes, with most sequence divergence confined to the N– and/or C-terminal tails depending on subtype (Fig. 1A,B; Fig. 2A,B). By contrast, the globular core domains, including residues involved in GTP binding and heterodimer formation, were nearly identical. Quantification of per-residue heterogeneity confirmed this pattern, with minimal divergence across the folded core and sharply increased variability toward the C-termini of both α– and β-tubulins (Fig. 1A,B; Fig. 2A,B).

**Figure 2.**
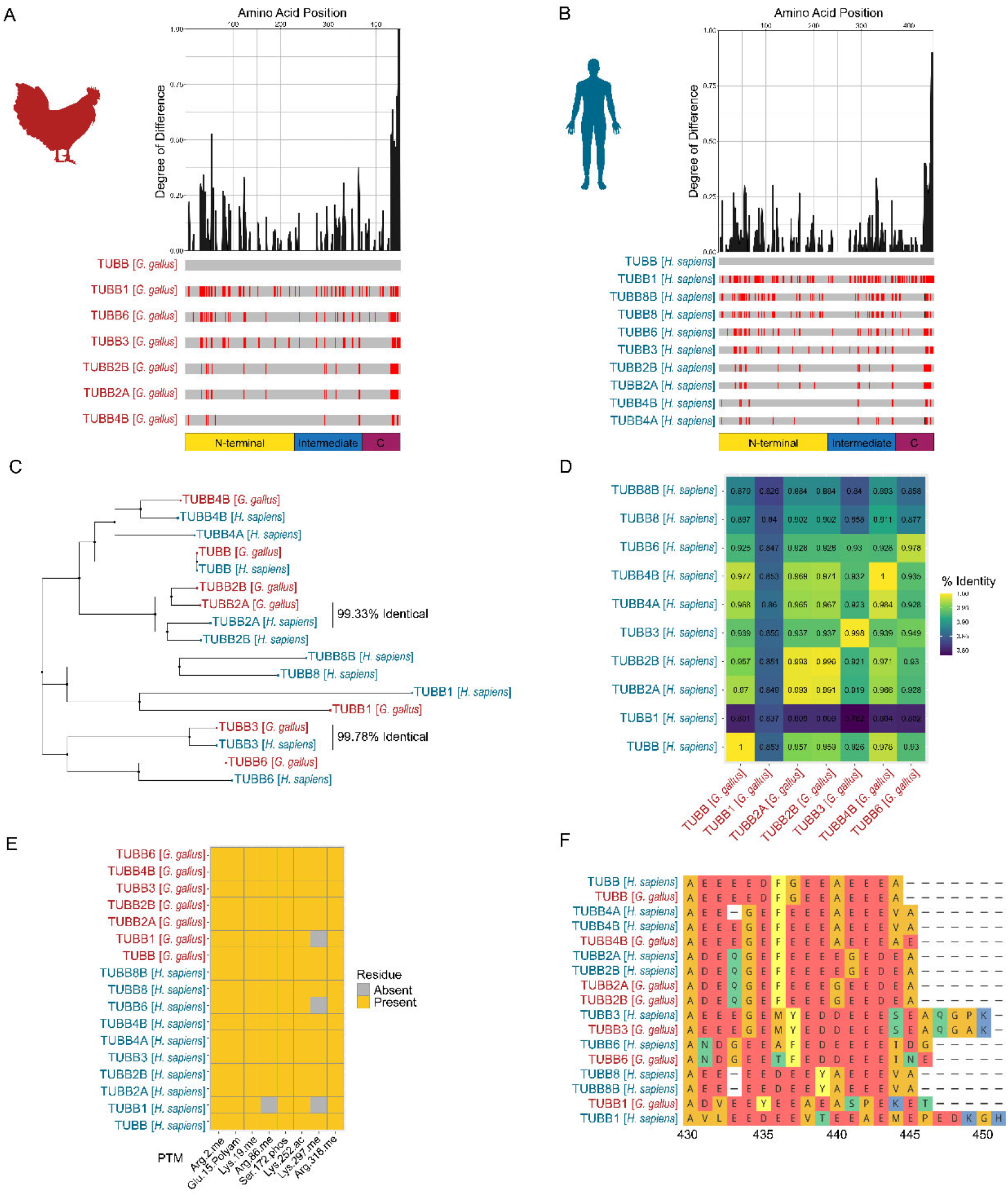
Sequence conservation and divergence of β-tubulin isotypes between chick and human. β-tubulin isotypes show strong cross-species conservation with selective C-terminal divergence, supporting structural conservation alongside potential regulatory specialization. (A,B) Multiple sequence alignments of β-tubulin isotypes from (A) *Gallus gallus* and (B) *Homo sapiens*. Amino acid mismatches relative to TUBA1A are shown in red, and gaps are shown in black. Plots above each alignment indicate per-residue heterogeneity, calculated as the complement of the conservation score averaged over three-residue bins. (C) Neighbor-joining phylogenetic tree of human and chick β-tubulin isotypes based on amino acid similarity. (D) Pairwise amino acid identity matrix for β-tubulin isotypes. (E) Distribution of canonical and predicted posttranslational modification sites across β-tubulin sequences. (F) Alignment of β-tubulin C-terminal tails. Human sequences are shown in blue and chick sequences in red.

Phylogenetic analysis further demonstrated that tubulin sequences clustered primarily by isotype rather than by species, indicating strong evolutionary constraint on isotype identity (Fig. 1C; Fig. 2C). With the exception of the closely related TUBB2A/B pair, chick and human orthologs consistently formed tight cross-species clades (Fig. 1C; Fig. 2C). Pairwise identity matrices reinforced these relationships, with orthologous chick-human pairs exhibiting the highest amino acid identity, often exceeding 90% (Fig. 1D; Fig. 2D).

Mapping canonical and predicted PTM sites onto the aligned sequences revealed broad conservation of putative modification sites within the globular domains, including residues associated with acetylation, methylation, and phosphorylation (Fig. 1E; Fig. 2E). By contrast, residues governing C-terminal detyrosination, polyglutamylation, and polyglycylation varied substantially among isotypes. Notably, the canonical C-terminal tyrosine required for α-tubulin tyrosination/detyrosination cycling was altered or absent in TUBA5/TUBA4A and the TUBAL3/TUBA8 group, suggesting isotype-specific constraints on tail-based PTM dynamics (Fig. 1E). Detailed alignment of C-terminal tails further revealed isotype-specific differences in tail length and charge distribution, with α-tubulins showing greater variability than β-tubulins (Fig. 1F; Fig. 2F).

Together, these analyses demonstrate that chick and human tubulin isotypes are highly conserved at structural and catalytic sites while diverging in their C-terminal tails, supporting the use of the chick as a model for investigating isotype-specific microtubule function and providing a molecular framework for interpreting tissue-level localization patterns.

### Spatiotemporal localization of TUBA1A during corneal development

To define the spatial and temporal distribution of select tubulin isotypes during corneal development, we performed IHC on chick embryos from embryonic day (E)3 through E14, encompassing corneal epithelial specification, periocular mesenchyme migration, stromal formation, and tissue maturation. We analyzed tubulin isotype differences through corrected total cell fluorescence (CTCF) to compare across cell types (Fig. S1,S2) and assessed apicobasal distribution profiles within the developing ocular monolayers (corneal epithelium, anterior lens epithelium, and corneal endothelium) (Fig. S3). We first characterized TUBA1A, and at E3, TUBA1A immunoreactivity was detected in the surface ectoderm, evenly across the presumptive corneal epithelium, as well as apically in the lens epithelium (Fig. 3A-D,Y,S2,S4). This pattern persisted in the corneal epithelium and lens through E4 and E5, during which lower fluorescence was identified in the periocular mesenchyme, which migrates and gives rise to the corneal endothelium and stroma (Fig. 3E-L,Y,S2,S4). By E6, when distinct corneal layers are established, TUBA1A was present throughout the epithelium, stroma, and endothelium, with a relatively even apicobasal distribution across the monolayers(Fig. 3M-P,S4).

**Figure 3.**
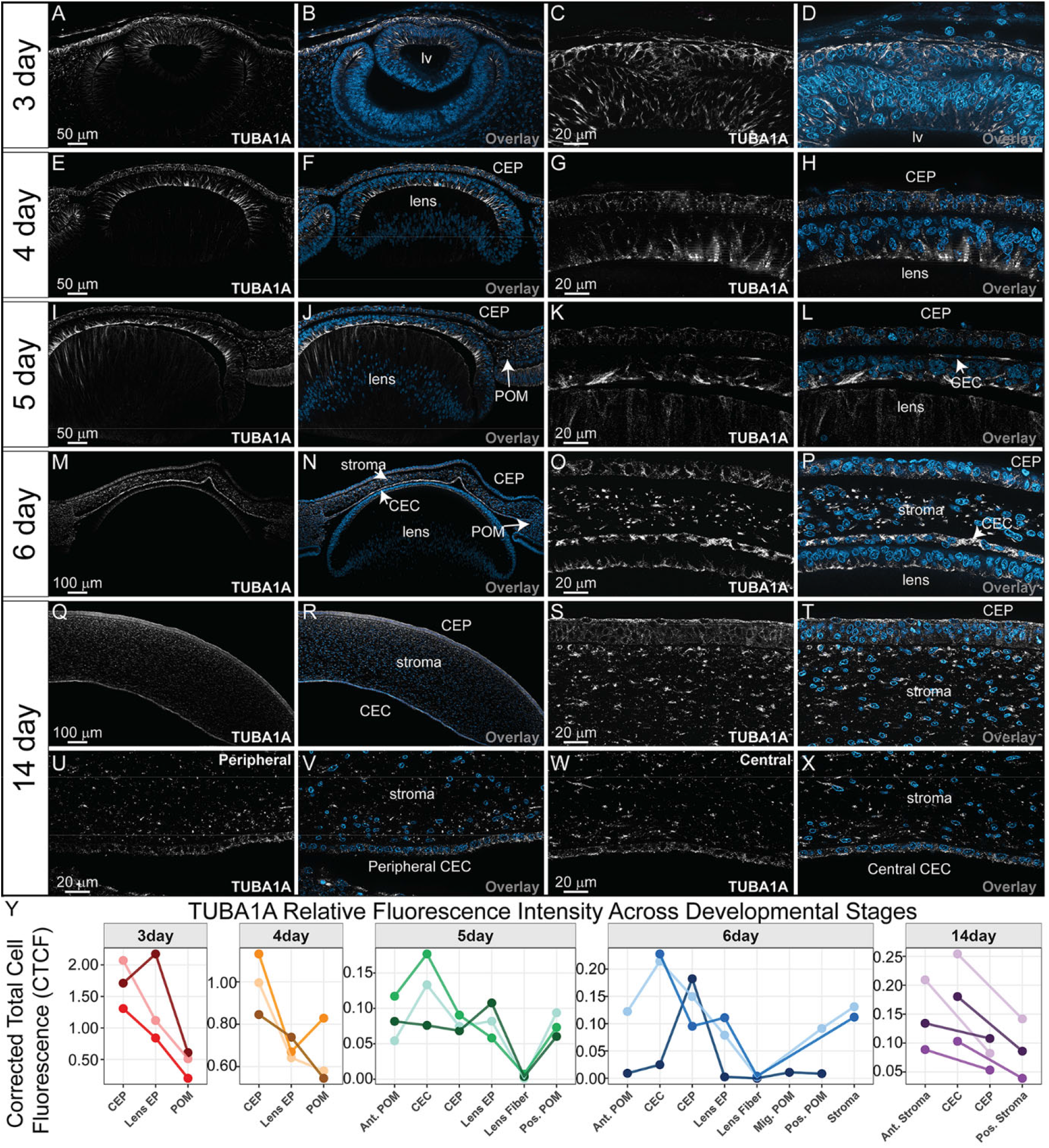
Spatiotemporal localization of TUBA1A during chick corneal development. TUBA1A exhibits broad expression throughout corneal development with stage-dependent redistribution and central enrichment in the mature endothelium. (A-X) Transverse cryosections after IHC in developing chick eyes at E3, E4, E5, E6, and E14 using antibodies against TUBA1A (white) and DAPI nuclear stain (blue). (A-D) E3 eye showing apical TUBA1A expression in the presumptive corneal epithelium. (E-H) E4 eye showing low, widespread expression during periocular mesenchyme migration. (I-L) E5 eye showing TUBA1A expression in the presumptive corneal endothelium. (M-P) E6 eye showing expression across corneal epithelium, stroma, and endothelium. (Q-X) E14 whole cornea showing strong epithelial and stromal expression. (U-X) Peripheral and central regions of the corneal endothelium at E14, showing central enrichment of TUBA1A. All sections are anterior to the top and posterior to the bottom. (Y) Relative fluorescence intensity of TUBA1A quantified using ImageJ across developmental stages and represented by corrected total cell fluorescence (CTCF). Individual data points represent the mean CTCF of three subarea measurements per cell type within a single biological replicate (N = 3). Connecting lines denote paired observations from the same biological replicate. Y axes are scaled independently to illustrate cell type CTCF variation. CEP, corneal epithelium; CEC, corneal endothelium; Lens EP, lens epithelium; POM, periocular mesenchyme; Ant., anterior; Pos., posterior; Mig., migratory.

In the maturing E14 cornea, TUBA1A exhibited robust expression in the apical stratifying corneal epithelium and stromal keratocytes (Fig. 3Q-X,S4). Within the corneal endothelium, robust TUBA1A signal was identified with an apically biased but more normally distributed fluorescence across the cell axis. (Fig. 3U-Y,S4). Across all developmental stages, TUBA1A was most expressed in the corneal epithelium or endothelium (Fig. 3A-Y,S2). These data demonstrate that TUBA1A is broadly expressed throughout corneal development but exhibits dynamic changes in subcellular localization and tissue distribution across time.

### Spatiotemporal localization of TUBA1B during corneal development

To compare the spatial distribution of closely related α-tubulin isotypes, we next examined the localization of TUBA1B across corneal development. The IHC analysis revealed that, although TUBA1B is broadly expressed, its localization pattern differs from that of TUBA1A. At E3, TUBA1B immunoreactivity was detected throughout the corneal epithelium, while a distinct band of enrichment was identified along the apical surface of the developing lens vesicle (Fig. 4A-D,Y,S5). This apical distribution in the anterior lens persisted at E4 and E5, during which TUBA1B remained broadly distributed across the developing corneal epithelium while also appearing in the apical presumptive corneal endothelium following periocular mesenchyme migration (Fig. 4E-L,Y,S5).

**Figure 4.**
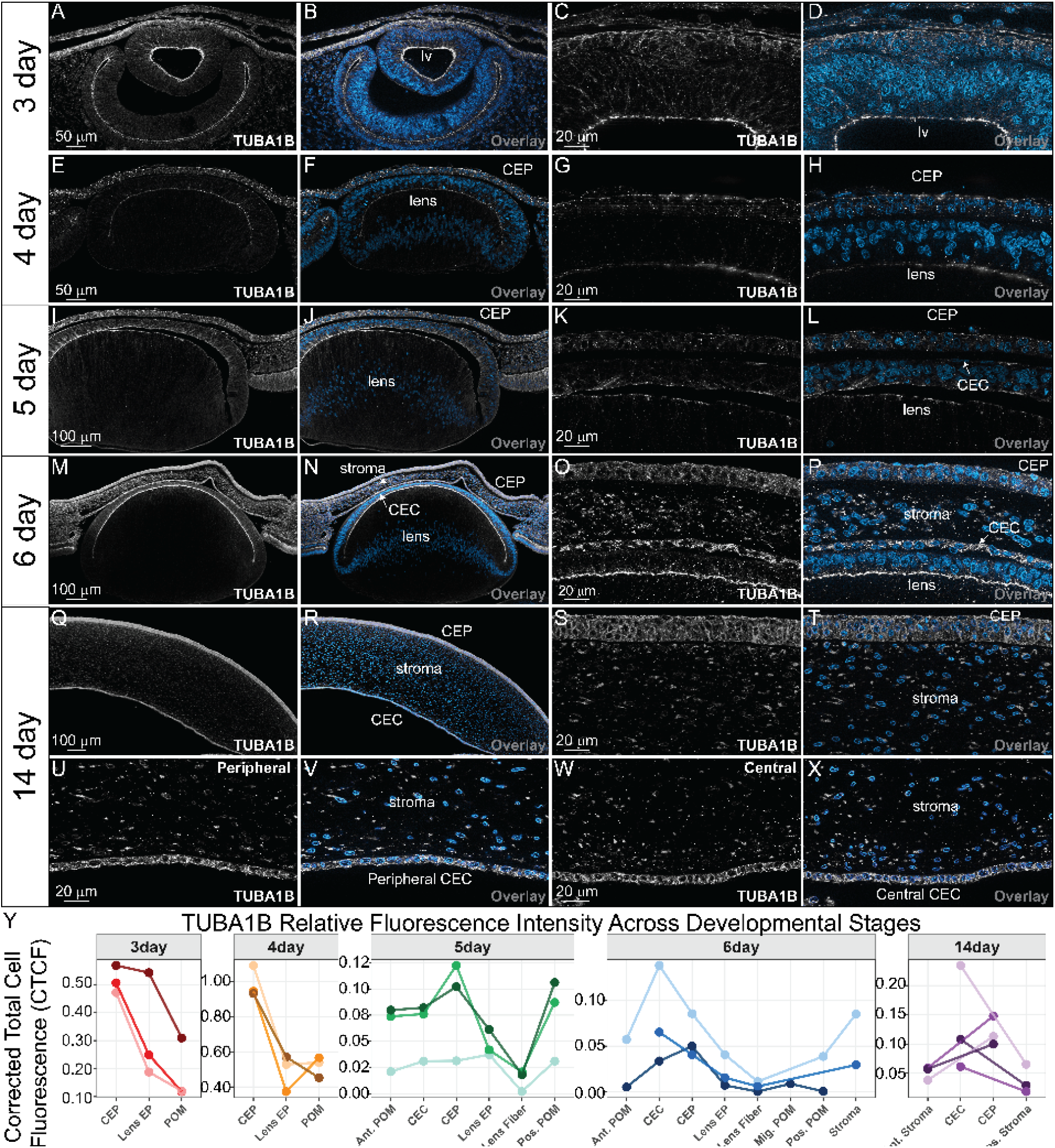
Spatiotemporal localization of TUBA1B during chick corneal development. TUBA1B is broadly and consistently expressed across corneal layers, displaying sustained epithelial enrichment and relatively uniform endothelial distribution. (A-X) Transverse sections of IHC in developing chick eyes at E3, E4, E5, E6, and E14 using antibodies against TUBA1B (white) and DAPI nuclear stain (blue). (A-D) E3 eye showing low-level, widespread expression in the surface ectoderm with apical enrichment in the presumptive lens epithelium. (E-H) E4 eye showing continued epithelial expression during periocular mesenchyme migration. (I-L) E5 eye showing expression in the presumptive corneal endothelium. (M-P) E6 eye showing cytoplasmic expression across corneal epithelium, stroma, and endothelium. (E-E‴) E14 whole cornea showing robust expression in stratifying epithelium and stroma. (Q-X) Central and peripheral regions of the corneal endothelium at E14 showing uniform endothelial expression. All sections are anterior to the top and posterior to the bottom. (Y) Relative fluorescence intensity of TUBA1B quantified via ImageJ across developmental stages and represented by corrected total cell fluorescence (CTCF). Individual data points represent the mean CTCF of three subarea measurements per cell type within a single biological replicate (N = 3). Connecting lines denote paired observations from the same biological replicate. Y axes are scaled independently to illustrate cell type CTCF variation. CEP, corneal epithelium; CEC, corneal endothelium; Lens EP, lens epithelium; POM, periocular mesenchyme; Ant., anterior; Pos., posterior; Mig., migratory.

By E6, when corneal layers are clearly established, TUBA1B exhibited a shift to a broadly cytoplasmic distribution throughout the corneal epithelium, stroma, and endothelium (Fig. 4M-P,Y,S5). Although the apical band remained visible in the anterior lens epithelium, more fluorescence was visible in the cytoplasm with a broader spread in fluorescence distribution across the apicobasal axis (Fig. 4M-P,Y,S5). In the mature E14 cornea, TUBA1B remained robustly expressed in the stratifying corneal epithelium, stromal keratocytes, and corneal endothelium (Fig. 4Q-X,Y,S5). In contrast to TUBA1A, TUBA1B displayed more uniform expression across the apicobasal axis of corneal endothelium and epithelium (Fig. 4U-X,Y,S5). These observations indicate that TUBA1B is a broadly expressed α-tubulin isotype throughout corneal development, with consistent enrichment in the epithelium, a comparatively uniform distribution within the endothelium, and a distinct apically polarized distribution in the lens epithelium.

### Spatiotemporal localization of TUBA5 (ortholog to human TUBA4A) during corneal development

To further compare α-tubulin isotype distribution during corneal morphogenesis, we examined the expression of TUBA5/TUBA4A across developmental stages. IHC analysis revealed that TUBA5/TUBA4A exhibits dynamic and spatially restricted localization patterns during corneal development, compared to the other ubiquitous α-tubulin isotypes. At E3, prior to periocular mesenchyme migration, TUBA5/TUBA4A expression was confined to the surface ectoderm, with strong enrichment at the apical surface of the presumptive corneal epithelium (Fig. 5A-D,AK,S2,S6). This apical localization was maintained at E4, with increased overall signal intensity in the corneal epithelium (Fig. 5E-H,AK,S6). By E5, following the first wave of periocular mesenchyme migration, TUBA5/TUBA4A was detected in both the corneal epithelium and the presumptive corneal endothelium at high levels (Fig. 5I-P,AK,S2,S6). Elevated signal was also observed in the anterior periocular mesenchyme contiguous with the forming endothelial layer (Fig. 5I-P [arrows],AK,S2,S6).

**Figure 5.**
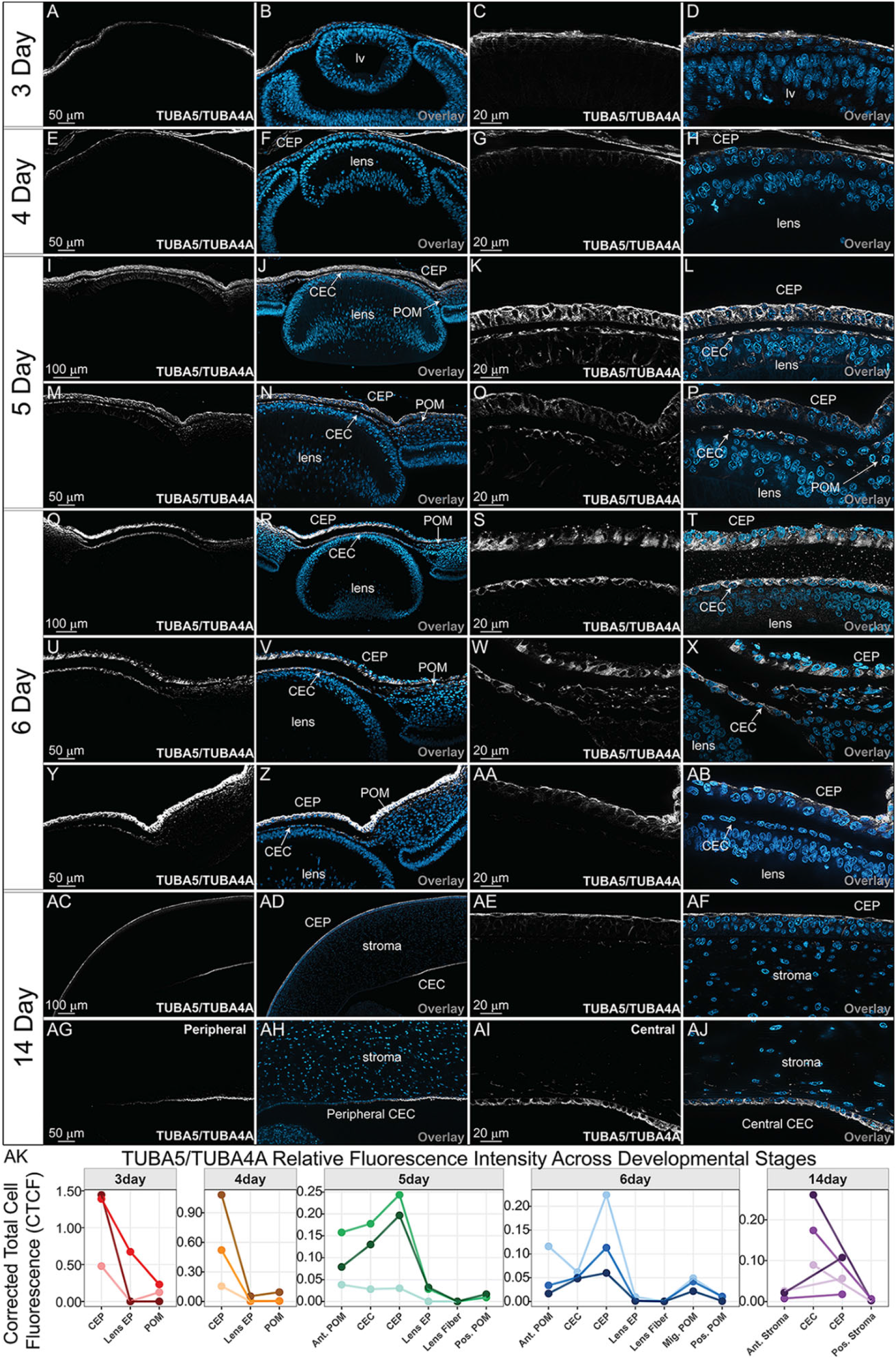
Dynamic and spatially restricted localization of TUBA5/TUBA4A during corneal development. TUBA5**/**TUBA4A shows dynamic, spatially restricted redistribution during corneal morphogenesis, with stage-specific shifts and central enrichment in the mature endothelium. (A-AJ) Transverse sections of IHC in developing chick eyes at E3, E4, E5, E6, and E14 using antibodies against TUBA5**/**TUBA4A (white) and DAPI nuclear stain (blue). (A-D) E3 eye showing apical expression in the presumptive corneal epithelium. (E-H) E4 eye showing increased apical epithelial expression. (I-P) E5 eye showing expression in the corneal epithelium, presumptive corneal endothelium, and anterior periocular mesenchyme. (Q-AB) E6 eye showing basal localization in epithelium and endothelium and expression in migrating stromal precursors. (AC-AJ) Central and peripheral regions of the corneal endothelium at E14 showing central enrichment and apical bias and expression restricted to the epithelium and endothelium. All sections are anterior to the top and posterior to the bottom. (Y) Relative fluorescence intensity of TUBA5/TUBA4A quantified via ImageJ across developmental stages and represented by corrected total cell fluorescence (CTCF). Individual data points represent the mean CTCF of three subarea measurements per cell type within a single biological replicate (N = 3). Connecting lines denote paired observations from the same biological replicate. Y axes are scaled independently to illustrate cell type CTCF variation. CEP, corneal epithelium; CEC, corneal endothelium; Lens EP, lens epithelium; POM, periocular mesenchyme; Ant., anterior; Pos., posterior; Mig., migratory.

At E6, when corneal layers are established, TUBA5/TUBA4A localization shifted toward a more basal distribution within both the corneal epithelium and endothelium (Fig. 5Q-AB,AK,S2,S6). Expression remained evident in the anterior periocular mesenchyme corresponding to the second wave of migratory stromal precursors (Fig. 5Y– AB,AK,S2). In the mature E14 cornea, TUBA5/TUBA4A expression was largely restricted to the corneal epithelium and endothelium, with minimal stromal signal (Fig. 5AC-AJ,AK,S2,S6). The TUBA5/TUBA4A distribution displayed an apical bias in the corneal epithelium and endothelium (Fig. 5AC-AJ,S6). These observations indicate that TUBA5/TUBA4A undergoes marked redistribution during corneal development and exhibits differential apicobasal patterning within the epithelial layers of the cornea.

Because TUBA5/TUBA4A showed the most spatially restricted and developmentally changing pattern among the α-tubulins examined, including enrichment in migratory periocular mesenchyme and the developing endothelium, we selected this isotype for higher-resolution analysis at E6. This stage allowed us to compare its organization in migratory mesenchymal cells and the epithelialized endothelial monolayer. We also examined TUBB4/TUBB3 co-distribution as an initial assessment of an accompanying β-tubulin isotype. To further resolve TUBA5/TUBA4A localization at cellular resolution and to identify possible overlap with putative β-tubulin isotypes during dynamic redistribution, we performed multiplexed IHC using antibodies for TUBA5/TUBA4A together with TUBB3 (Fig. 6A-H). At E6, we identified overlap between the two isotypes in the corneal epithelium and forming endothelium in the central cornea (Fig. 6A-D). While TUBA5/TUBA4A was localized to the peripheral corneal endothelium and migratory periocular mesenchyme, the TUBB3 signal was sparse in these cells (Fig. 5O-AB, 6E-H). Additional confocal analyses on flat-mounted cornea to better visualize subcellular localization of TUBA5/TUBA4A at E6 revealed a diffuse cytosolic and peri-membranous localization of TUBA5/TUBA4A across the epithelialized corneal endothelial monolayer when visualized together with the junctional marker N-cadherin (NCAD) and filamentous (F-)actin (Fig. 6I-M’). In dividing endothelial cells, TUBA5/TUBA4A signal was enriched in microtubule structures associated with separating chromatin (Fig. 6I,I’, M,M’). F-actin was primarily localized to the cortex of the endothelium (Fig. 6J,J’,M,M’). NCAD was localized in and around the membrane, with punctate staining in the cytoplasm (Fig. 6K,K’,M,M’). In the migratory periocular mesenchyme, TUBA5/TUBA4A labeled prominent microtubule bundles of variable diameter (Fig. 6N,N’,R,R’), which were particularly evident at the leading edge of the migratory population (Fig. 6N, N’). The F-actin staining in these cells appeared more diffuse and included cytosolic puncta and filopodial projection (Fig. 6O,O’,R,R’). NCAD staining was almost absent in the leading edge of the migratory cells (Fig. 6P,P’). Together, these data demonstrate that TUBA5/TUBA4A exhibits dynamic subcellular localization patterns in both epithelialized endothelial cells and migratory mesenchymal populations during corneal development.

**Figure 6.**
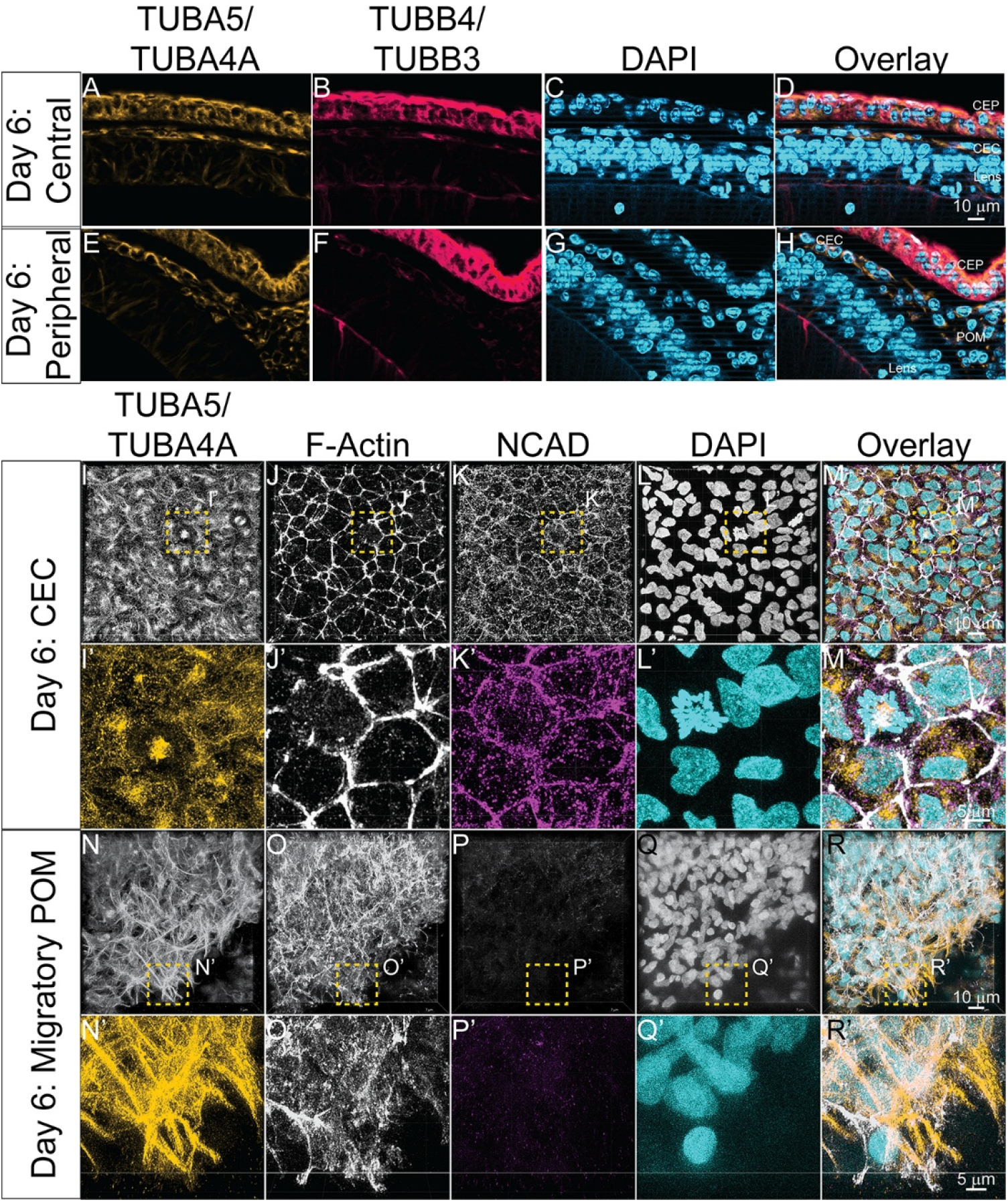
TUBA5/TUBA4A dynamics during early corneal development. Flat mount imaging reveals contextdependent TUBA5/TUBA4A organization, with diffuse peri-membranous distribution in endothelial cells and prominent bundled microtubules in migratory mesenchyme. (A-H) Transverse sections of IHC in developing chick eyes at E6 in the central and peripheral cornea using antibodies against TUBA5/TUBA4A (yellow), TUBB3 (red), and DAPI nuclear stain (blue). (I-R’) IHC of corneal flat mounts at E6 using antibodies against TUBA5/TUBA4A (yellow), F-actin stain (white), NCAD (magenta), and DAPI (cyan). (I-M) E6 corneal endothelial maximum intensity projections (MIP) show discrete cortical F-actin and membranous NCAD with diffusive distribution of small diameter TUBA5/TUBA4A microtubules. (I’-M’) Zoomed MIP shows membrane NCAD, cortical actin, diffuse perimembranous and centrosomal TUBA4A staining in corneal endothelia. (N-R) E6 migratory periocular mesenchyme MIP shows diffuse F-actin, minimal NCAD staining, with discrete large diameter TUBA5/TUBA4A microtubules biased towards the leading edge. (N’-R’) Zoomed MIP highlights predominantly diffuse and minor punctate NCAD staining in addition to large TUBA5/TUBA4A microtubules. N= 9 corneas assessed for 6-day flat mounts.

### Spatiotemporal localization of TUBB1 (ortholog to human TUBB2A) during corneal development

To assess β-tubulin isotype distribution during corneal morphogenesis, we examined the localization of TUBB2A across developmental stages. TUBB1/TUBB2A exhibited broad expression but with notable temporal changes in epithelial localization. At E3, TUBB1/TUBB2A immunoreactivity was detected robustly throughout the surface ectoderm and the anterior lens epithelium, with modest enrichment at the apical surface of the presumptive corneal epithelium (Fig. 7A-D,Y,S2,S7). This pattern was maintained at E4, with continued expression in the corneal epithelium and periocular mesenchyme (Fig. 7E-H,Y,S7). By E5, following establishment of the presumptive corneal endothelium, TUBB1/TUBB2A polarized apical fluorescence was decreased in addition to increasing generalized apicobasal expression within endothelial cells (Fig. 7I-L,Y,S2,S7).

**Figure 7.**
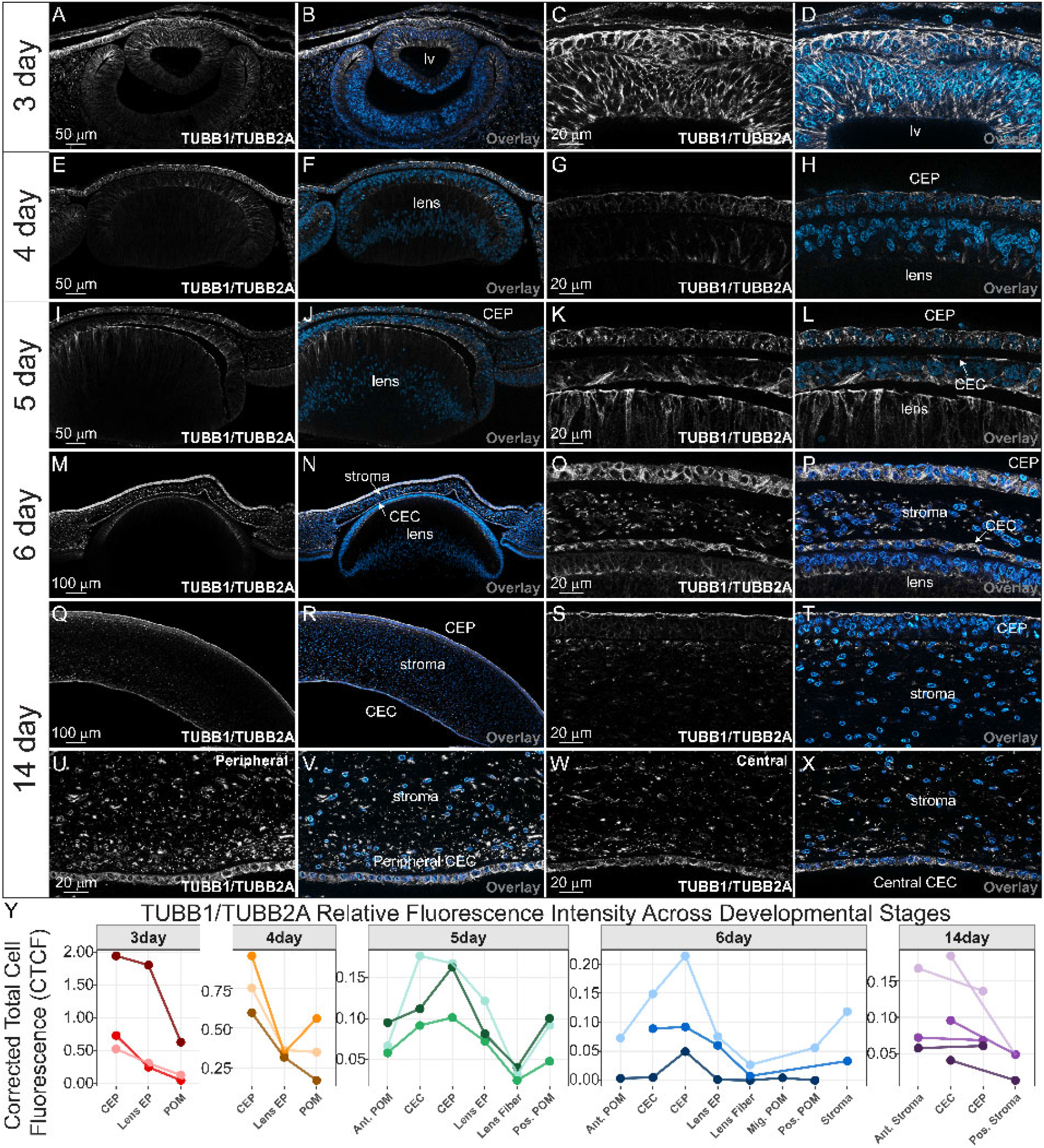
Developmental redistribution of TUBB1/TUBB2A in corneal tissues. TUBB1/TUBB2A undergoes developmental redistribution across corneal tissues, transitioning from broad early epithelial expression to polarized enrichment in mature layers. (A-X) Transverse cryosection of IHC in developing eye after 3, 4, 5, 6 and 14 days of incubation with antibodies against TUBB1/TUBB2A protein (white) and DAPI. (A-D) E3 eye showing broad epithelial expression with apical enrichment. (E-H) E4 eye showing continued epithelial and periocular mesenchyme expression. (I-L) E5 eye showing reduced epithelial apical signal and increased endothelial expression. (M-P) E6 eye showing cytoplasmic expression across corneal layers. (Q-T) E14 whole cornea showing apical epithelial expression and stromal localization. (U-X) Central and peripheral regions of the corneal endothelium at E14 show central enrichment and apical bias. All sections are anterior to the top and posterior to the bottom. (Y) Relative fluorescence intensity of TUBB2A quantified via ImageJ across developmental stages and represented by corrected total cell fluorescence (CTCF). Individual data points represent the mean CTCF of three subarea measurements per cell type within a single biological replicate (N = 3). Connecting lines denote paired observations from the same biological replicate. Y axes are scaled independently to illustrate cell type CTCF variation. CEP, corneal epithelium; CEC, corneal endothelium; Lens EP, lens epithelium; POM, periocular mesenchyme; Ant., anterior; Pos., posterior; Mig., migratory.

At E6, TUBB1/TUBB2A remained broadly expressed across the corneal epithelium, stroma, and endothelium, with a predominantly cytoplasmic distribution and reduced apical bias across all ocular monolayers (Fig. 7M– P,Y,S2,S7). In the mature E14 cornea, TUBB1/TUBB2A expression increased at the apical surface of the stratified corneal epithelium and remained robust within anterior stromal keratocytes (Fig. 7Q-X,Y,S2,S7). Within the corneal endothelium, TUBB1/TUBB2A was expressed throughout the tissue, with an apical bias and a higher fluorescence than the posterior stromal cells (Fig. 7U-X,Y,S2,S7). These findings indicate that TUBB1/TUBB2A undergoes developmental redistribution within the corneal and lens epithelium while maintaining broad expression in the corneal endothelium.

### Spatiotemporal localization of TUBB4 (ortholog to human TUBB3) during corneal development

Finally, we examined the expression of TUBB4/TUBB3, a neuron-associated β-tubulin isotype, to determine its distribution within corneal tissues. TUBB4/TUBB3 displayed a highly polarized and developmentally persistent localization pattern. At E3, TUBB4/TUBB3 immunoreactivity was detected most highly in the surface ectoderm, corresponding to the presumptive corneal epithelium (Fig. 8A-D,Y,S2). The TUBB4/TUBB3 signal maintained an apical skew at E4 and E5, with low to moderate expression observed in the migrating periocular mesenchyme and developing corneal endothelium at E5 (Fig. 8E-L,S2,S8). At E6, TUBB4/TUBB3 was detected in both the corneal epithelium and endothelium, but their CTCF values were not significantly different in the within-stage analysis (Fig. 8Y,S2).

**Figure 8.**
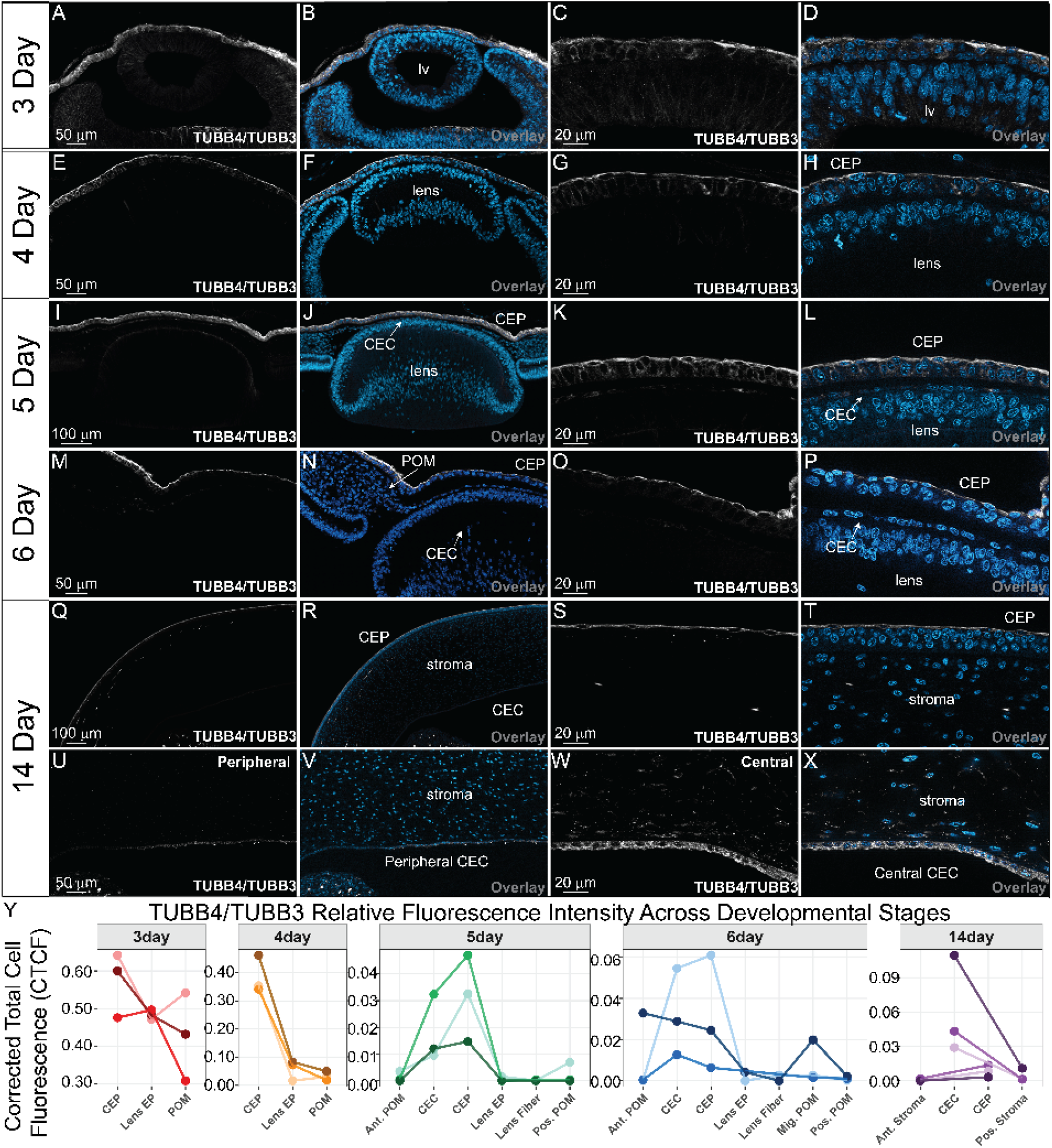
Polarized expression of TUBB4/TUBB3 in corneal epithelium, endothelium, and stromal nerves. TUBB4/TUBB3 displays polarized apical epithelial localization throughout development and selectively labels stromal nerves in the mature cornea. (A-X) Transverse sections of IHC in developing chick eyes at E3, E4, E5, E6, and E14 using antibodies against TUBB4/TUBB3 (white) and DAPI nuclear stain (blue). (A-D) E3 eye showing apical epithelial expression. (E-H) E4 eye showing sustained apical localization. (I-L) E5 eye showing apical epithelial expression and limited endothelial signal. (M-P) E6 eye showing similar epithelial localization with some corneal endothelial expression. (Q-X) E14 whole cornea showing strong apical epithelial expression and stromal nerve labeling. (Q-X) Central and peripheral regions of the corneal endothelium at E14 show central enrichment and apical bias. All sections are anterior to the top and posterior to the bottom. (Y) Relative fluorescence intensity of TUBB4/TUBB3 quantified via ImageJ across developmental stages and represented by corrected total cell fluorescence (CTCF). Individual data points represent the mean CTCF of three subarea measurements per cell type within a single biological replicate (N = 3). Connecting lines denote paired observations from the same biological replicate. Y axes are scaled independently to illustrate cell type CTCF variation. CEP, corneal epithelium; CEC, corneal endothelium; Lens EP, lens epithelium; POM, periocular mesenchyme; Ant., anterior; Pos., posterior; Mig., migratory.

In the mature E14 cornea, TUBB4/TUBB3 was most enriched at the apical surface of the stratified corneal epithelium and within the corneal endothelium (Fig. 8Q-X,Y,S2,S8). In addition, prominent TUBB4/TUBB3-positive tracks were observed within the anterior stroma, consistent with developing corneal nerves (Fig. 8Q-X). Analysis of endothelial localization revealed an apical bias (Fig. 8U-Y,S8). These results demonstrate that TUBB4/TUBB3 marks apical microtubule populations in the corneal epithelium throughout development and becomes increasingly associated with central endothelial regions and stromal innervation during corneal maturation.

## Discussion

We combine chick-human tubulin sequence comparisons with quantitative developmental imaging to map five tubulin isotypes during corneal morphogenesis (Fig. 9). Within-stage fluorescence measurements and normalized apicobasal profiles reveal differences in tissue enrichment and developmental localization, while E6 co-staining further defines TUBA5/TUBA4A distribution relative to TUBB4/TUBB3 and epithelial organization. This atlas identifies specific stages and cell populations for testing how tubulin composition contributes to corneal development.

**Figure 9.**
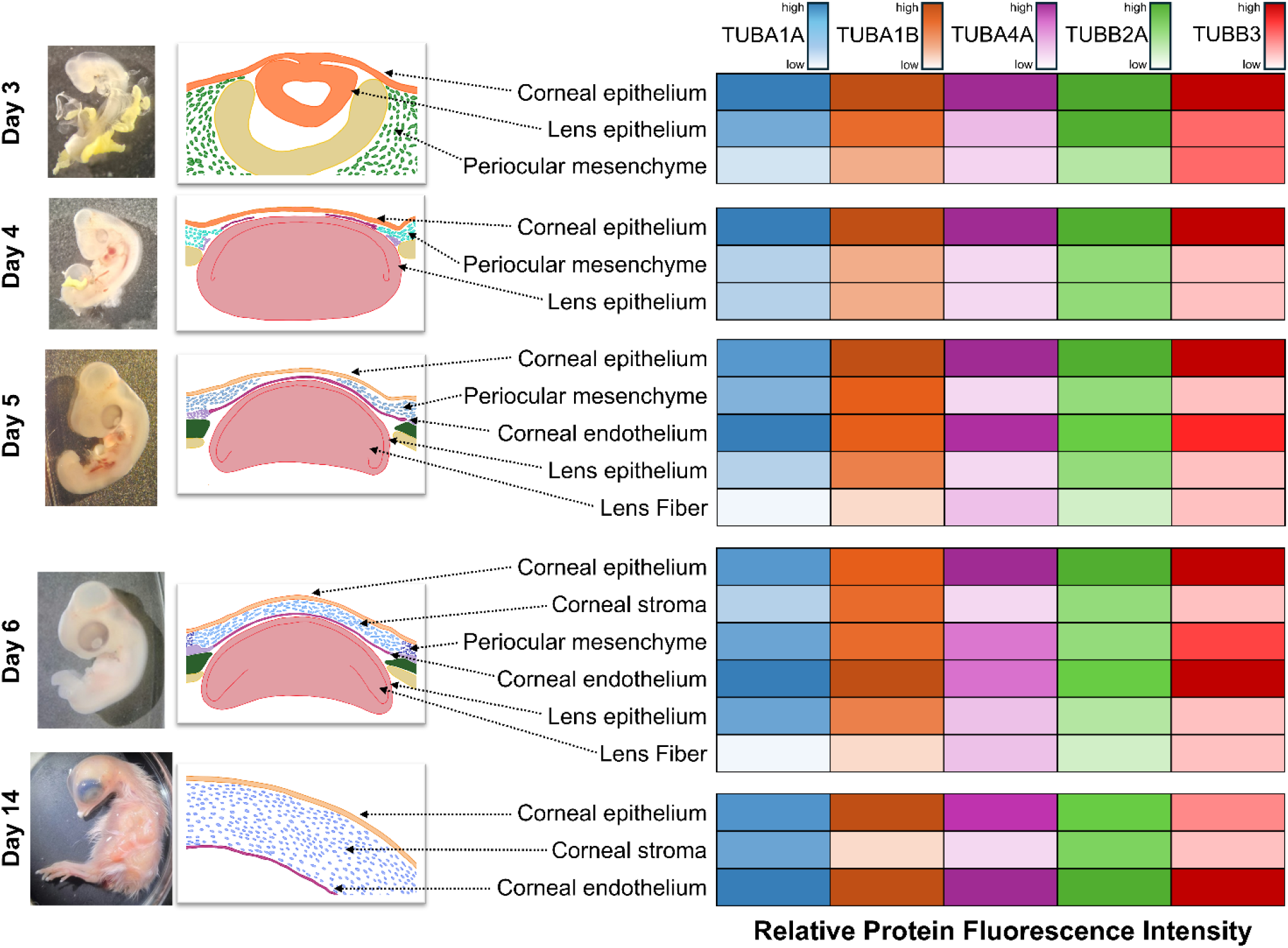
Graphical illustration of tubulin isotype spatiotemporal distribution in the developing chick cornea. Visualization show cases relative fluorescence intensity of tubulin isotypes across major developing corneal and corneal adjacent cell types in the chick embryo. Darker color indicates higher fluorescence while lower saturation indicates lower fluorescence. Raw images of chick embryos shown on the left along with illustrations of corneal cross sections based on select tissue sections.

Our comparative sequence analyses revealed that both α– and β-tubulin families cluster by isotype rather than by species, underscoring strong evolutionary constraint on isotype identity (Fig. 1, 2). Conserved core domains provide a basis for cross-species comparison, while divergent tails identify candidate sites of regulatory specialization (Vemu, Atherton et al. 2016). Instead, sequence divergence is concentrated within the N– and/or C-terminal tails, consistent with nucleotide binding domains (Yon, Ha et al. 2025) and the established role of C-termini as platforms for PTMs and interactions with motor proteins and MAPs (Gadadhar, Bodakuntla et al. 2017, Janke and Magiera 2020, Bao, Dorig et al. 2023). Mapping of canonical and predicted PTM sites further supports this view, as modification sites within the globular core are largely conserved, whereas residues governing detyrosination, polyglutamylation, and polyglycylation vary across isotypes. These differences raise the possibility that isotype composition may help determine the intrinsic properties of microtubules and which PTMs can be acquired and how the resulting microtubule is interpreted by motors and MAPs.

Isotype-selective PTM regulation has been demonstrated in several experimental systems but remains incompletely characterized in developing vertebrate tissues. In *Drosophila* ovaries, αTub84B and αTub84D are glutamylated, whereas the ovary-enriched αTub67C is not, demonstrating that closely related α-tubulin isotypes can be differentially modified *in vivo* (Bao, Dorig et al. 2023). In *C. elegans*, sequence variations among C-terminal tails similarly alters the availability of glutamate residues that can serve as substrates for polyglutamylation and polyglycylation, providing a mechanism by which isotype identity could constrain the PTM repertoire (Lu and Zheng 2022). Vertebrate evidence also links isotype-specific PTMs to MAP interactions. For example, preventing C-terminal polyglutamylation of mouse TUBA4A reduces the association of Tau and MAP2 with neuronal microtubules (Hausrat, Janiesch et al. 2022). These findings establish that PTM-dependent regulation of a defined tubulin isotype can influence MAP binding *in vivo*, although corresponding preferences in developing corneal tissues remain unknown. Taken together, prior studies support a model in which the tubulin code is combinatorial; genetically encoded isotype diversity can influence the subsequent acquisition of PTMs, thereby generating microtubule populations with distinct biochemical properties.

PTMs can in turn alter the recruitment or activity of specific microtubule-interacting proteins. Tyrosination promotes interactions with several CAP-Gly-containing proteins and kinesin-13 family members, while detyrosination can alter kinesin-dependent transport and polyglutamylation can regulate kinesin motility and the activity of microtubule-severing enzymes like spastin and katanin (Song and Brady 2015, Janke and Magiera 2020). Compelling evidence for interplay between isotype identity and PTM state comes from reconstituted microtubules with defined C-terminal tail composition. Kinesin-1 has a reduced run length on microtubules containing the TUBB3 C-terminal tail compared to the TUBB2 tail, which can be attributed to the terminal lysine unique to TUBB3. However, addition of polyglutamate chains to the paired TUBA1A C-terminal tail can restore kinesin-1 processivity on TUBB3-containing microtubules (Sirajuddin, Rice et al. 2014). Therefore, the functional properties of a microtubule cannot necessarily be predicted from either isotype identity or PTM state alone. Rather, α/β-isotype composition and PTMs act combinatorially to regulate motor behavior and further analysis of tubulin PTMs during corneal morphogenesis is necessary to identify if these aspects of the tubulin code are relevant during morphogenesis.

The framework noted above is relevant to the developmental patterns described here. The changing abundance and localization of α– and β-tubulin isotypes during corneal morphogenesis could establish microtubule populations with different capacities for modification and different affinities for motors and MAPs. However, the extent to which individual vertebrate tubulin isotypes are preferentially modified *in vivo* remains incompletely defined. Our sequence-based PTM analysis identifies potential modification sites rather than directly measuring PTM occupancy. Determining whether the spatially restricted isotype populations identified here acquire distinct PTM signatures, and whether these combinations selectively recruit motors or MAPs, represents an important direction for future studies of corneal morphogenesis.

TUBA5/TUBA4A exhibits the most dynamic and spatially restricted pattern among the α-tubulins examined (Fig. 5,6). Its early apical enrichment in the presumptive corneal epithelium, transient expression in migrating periocular mesenchyme, and later restriction to epithelial and endothelial compartments, coupled with pronounced gradients in the endothelium, point to tightly regulated deployment during morphogenesis. TUBA4A is unusual among vertebrate α-tubulins because it is genetically encoded without the canonical C-terminal tyrosine. Recent biochemical studies further demonstrate that human TUBA4A is a relatively poor substrate for tubulin tyrosine ligase compared with detyrosinated TUBA1B, and that transferring the TUBA4A CTT to the TUBA1B core is sufficient to confer reduced tyrosination (Fu, Yan et al. 2023). Thus, incorporation of TUBA5/TUBA4A may provide a direct mechanism for generating microtubule populations with a distinct tyrosination state rather than simply serving as a substrate for differential post-translational processing. Differences in TUBA5/TUBA4A microtubule morphology were identified between epithelialized corneal endothelia and the migratory mesenchyme suggestive of overlapping structural roles in both cell states. The localization of TUBA5/TUBA4A at the leading edge of migratory corneal progenitors and within the maturing endothelium is consistent with roles in stabilizing microtubule arrays during collective cell migration and in maintaining long-lived epithelial monolayers (Etienne-Manneville 2013, Bar, Popp et al. 2022). Given its importance at this late stage of corneal development, future work should focus on the role of TUBA5/TUBA4A in development and disease.

Distinct tubulin isotypes can co-assemble into mixed microtubules, as demonstrated using chicken brain and erythrocyte tubulin preparations (Baker, Rothwell et al. 1990). However, our tissue-level localization profiles and E6 co-staining do not determine the relative proportions or spatial arrangement of isotypes within individual corneal microtubules. Whether epithelial, endothelial, and mesenchymal cells contain microtubules with different degrees of isotype mixing therefore remains unresolved. Future filament-resolved imaging could address this question, while controlled isotype replacement could test the contribution of cell-type-specific composition to microtubule dynamics and tissue organization. Such experiments should preserve total tubulin abundance, assess compensatory changes in other isotypes, and compare rescue by the native isotype with rescue by an alternative isotype or a C-terminal-tail variant. These approaches would distinguish general tubulin requirements from isotype-dependent effects on polarity, migration, and endothelial monolayer formation.

In contrast to α-tubulins, β-tubulin isotypes displayed sharper spatial biases. TUBB1/TUBB2A is broadly expressed across corneal tissues but undergoes developmental redistribution within the epithelium, transitioning from early apical enrichment to a more cytoplasmic pattern and later re-establishing apical localization during epithelial stratification (Fig. 7). Within the corneal endothelium, TUBB1/TUBB2A remains robustly expressed and develops an apically biased gradient during maturation. TUBB4/TUBB3, although classically associated with neuronal microtubules (Katsetos, Herman et al. 2003, Jouhilahti, Peltonen et al. 2008), has also previously been identified in non-neuronal migratory neural crest cell populations (Chacon and Rogers 2019). TUBB4/TUBB3 shows persistent apical enrichment in the corneal epithelium throughout development and becomes increasingly prominent in central endothelial regions and stromal nerve tracks at later stages (Fig. 8). These patterns provide a framework for future studies examining whether β-isotypes preferentially associate with dynamic or polarized microtubule populations.

Investigating tubulin isotype apicobasal distribution profiles across the corneal epithelium and endothelium highlights an increasing apical skew with development (Fig. S1-S8). We noted that TUBA5/TUBA4A and TUBB3 share the most similar profiles in the corneal epithelium, with striking apical polarization (Fig. 6,S6,S8). A similar trend is noted in the corneal endothelium, although most tubulins seem to share overlapping profiles. These overlapping profiles identify candidate co-distribution patterns but do not establish preferential α/β pairing, co-assembly within individual microtubules, or shared functional output. This may highlight the importance of specialized subapical microtubule arrays in epithelial layer polarity. In the differentiated corneal epithelial layer, the apical surface consists of a multitude of microvilli and microplicae, critical to ocular surface health (Sharkova, Chow et al. 2023, Troisi, Del Prete et al. 2024). The corneal endothelium relies on long-lived microtubules to sustain vesicle trafficking and junctional integrity (Bonanno 2012). Additionally, primary cilia develop on the apical surface of corneal endothelial cells during embryogenesis and wound healing, linking microtubules to these apical specializations in health (Blitzer, Panagis et al. 2011). Differential isotype composition may contribute to these regional functional demands in these layers.

Importantly, the developmental patterns described here intersect with emerging links between microtubule dysregulation and human corneal disease. Altered tubulin expression and microtubule remodeling have been implicated in the pathogenesis of Fuchs’ endothelial corneal dystrophy and other corneal disorders, and mutations in multiple tubulin genes are now associated with ocular phenotypes (Tischfield, Cederquist et al. 2011, Ong Tone, Kocaba et al. 2021). The conservation of tubulin isotypes and their developmental deployment in the chick cornea underscores the utility of this model for probing how isotype composition and PTM states contribute to corneal homeostasis and disease susceptibility.

The atlas provides a developmental reference for testing how isotype composition and PTM state contribute to corneal organization and disease.

## MATERIALS AND METHODS

### Sequence Analysis

All FASTA protein sequences were collected from the NCBI RefSeq database and aligned within the alpha– and beta-tubulin subtypes using the *msa* package in R. All subsequent computations were performed in R using these aligned sequences. Dendrograms were created using a neighbor-joining tree estimation algorithm using the *ape* package and further visualized using the *ggtree* package. Multiple sequence alignments (MSA) of the C-terminal tails was performed using the *ggmsa* package, while *seqvisr* was used to visualize gaps and mismatches across the entire alignment. In conjunction, heterogeneity across the sequence was visualized by calculating the degree of difference. This was defined by calculating a conservation score for each residue based on amino acid similarity, scaled from 0 to 1, then taking its complement such that a value of 0 indicated perfect conservation and 1 representing maximal diversity. To visualize overarching patterns in the sequences, values were averaged over bins of 3 residues. All sequence alignments, phylogenetic analyses, and image datasets generated in this study are available as a resource for comparative studies of tubulin isotype organization. All fluorescence raw data is available in a public github repository: https://github.com/luckycharms10/Rogers-Lab-Tubulins-Developing-Chick-Cornea.

### Chicken Eggs and Incubation

Fertilized chicken eggs were obtained from the UC Davis Hopkins Avian Facility at the University of California, Davis and incubated at 37 °C for approximately 3, 4, 5, 6, and 14 days.

### Dissection and Fixation

For 3 and 4-day samples, whole embryos were removed and fixed in 2% trichloroacetic acid (TCA) for 30 minutes at room temperature. For 5-day samples, the whole head was removed and fixed in 2% TCA for 40 minutes at room temperature. For 6-day samples, whole eyes were dissected and fixed in 2% TCA for 40 minutes at room temperature. For 14-day samples, only the anterior section of the eye was dissected along the limbal margin. The eyelid and lens were removed before the samples were fixed in 2% TCA for 30 minutes at room temperature. Corneal samples for flat mount images (Fig. 6) were obtained via dissection of 6-day embryos and fixed in 4% paraformaldehyde for 40 minutes.

### Immunohistochemistry

Immunohistochemistry (IHC) was performed based on a modified version of the protocol used previously (Echeverria, Leathers et al. 2025), using the primary antibodies listed in Table 3. After dissection and fixation, all samples were washed in 1X PBS (137 mM NaCl, 11.9 mM phosphates, 2.7 mM KCl; pH 7.4) containing 0.1% Triton X-100 (PBST). Samples were incubated in a blocking buffer made up of PBST with 10% donkey serum for 1-7 days at 4 °C. Primary antibodies were diluted in blocking buffer to the indicated dilution and incubated with samples for approximately 72 hours at 4 °C. After primary incubation, samples were washed with PBST then incubated with AlexaFluor secondary antibodies diluted in blocking buffer (1:500) for up to 24 hours at 4 °C. Samples were then washed in PBST and were post-fixed in 4% PFA for 1 hour at room temperature. Tissue samples were also incubated with a DAPI stain (1:500) to mark nuclei. For flat mount imaging (Fig. 6), 6-day dissected corneas subject to IHC as previously described were mounted on a slide with Fluoromount-G, cover slipped, and imaged.

**Table 3.** Antibodies Used in Study.

| Antibody Target | Antigen | Dilution | Species Reactivity | Host/Isotype | Source |
| --- | --- | --- | --- | --- | --- |
| Tubulin alpha-1A chain (TUBA1A) | Chicken native chick brain microtubules | 1:250 | human, mouse, gerbil, chicken, rat | Mouse/IgG1 | MilliporeSigma |
| Tubulin alpha-1B chain (TUBA1B) | Recombinant human Tubulin-Alpha protein | 1:250 | human, mouse, rat, rabbit, monkey, chicken, zebrafish, hamster, goat, p. lividus | Rabbit/IgG | Proteintech |
| Tubulin alpha-4A chain (TUBA4A) | Full-length protein | 1:250 | human | Mouse/IgG2b | ThermoFisher |
| Tubulin beta-2A chain (TUBB2A) | Full-length recombinant protein | 1:250 | human, mouse, rat | Mouse/IgG2a | Abnova |
| Tubulin beta-3 chain (TUBB3) | Peptide | 1:250 | human, mouse, rat, pig, rabbit, chicken | Mouse/IgG1 | Proteintech |
| Neuron-specific beta -III Tubulin Antibody | Rat brain-derived microtubules | 1:250 | Human, Mouse, Rat, Porcine, Amphibian, Avian, Avian, Primate, Transgenic Mouse, Transgenic Rat, Xenograft | Mouse/IgG2a | R&D Systems |
| N-cadherin (NCAD) | Amino acids 308-597 | 1:5 | Chick, human, mouse, *xenopus* | Rat/IgG2a | DSHB |

After IHC, samples for cross-sectional imaging were prepared for cryosectioning by washing with PBST, then subsequently incubated in 5% sucrose for 12-24 hours at 4 °C, followed by 15% sucrose for 12-24 hours at 4 °C. Samples were then embedded in gelatin and frozen with liquid nitrogen for cryosectioning. Whole mount images were taken of all samples following post-fixation, and transverse sections were taken after cryosectioning using a Thermo Scientific HM525 NX Cryostat.

### Fluorescence Imaging and Image Processing

All transverse section images were taken using a Zeiss Axio Imager.M2 with Apotome capability at 10X, 20X, or 63X magnification. Flat mount images (Fig. 6) were acquired on a Leica SP8 with STED capability 3X confocal microscope controlled by Leica LAS X software using a 100x/1.4 oil (HC PL APO CS2) object. Z-stacks were acquired with a 0.3-micron step size. All images were deconvolved on-the-fly with Huygens Batch Feeder version 25.10 using the “Standard” strategy (Scientific Volume Imaging, The Netherlands, http://svi.nl). Maximum Intensity Projections (MIP) were generated with Zeiss Zen optical processing software, Imaris (version 10). Imaris and Adobe Photoshop were used for fluorescence image processing. Images shown in manuscript are representative from 3-9 corneas per marker per stage.

### Image analysis and fluorescence quantification

Image processing and spatial quantification of fluorescence intensity was performed on ImageJ (Schneider, Rasband et al. 2012). For each isotype, three tissue sections from separate biological replicates were utilized (N=3).

For fluorescence intensity comparisons across cell types (presented in Fig. 3Y, 4Y, 5AK, 7Y, 8Y; Supplementary S1, S2), distinct cell type regions of interest (ROI) including the corneal epithelium (CEP), corneal endothelium (CEC), lens epithelium (Lens EP), periocular mesenchyme (POM) populations and stromal populations were manually delineated by stage in ImageJ. To account for localized within-cell type variability, three independent sub area ROIs were measured per cell type per biological replicate. For each ROI, the area and mean fluorescence were measured. The integrated density (IntDen) was computed as a product of the area and the mean intensity. To isolate biological signal from technical noise, the corrected total cell fluorescence (CTCF) was calculated for each ROI using the formula: CTCF = Mean IntDen – (Area x Mean Background Intensity). The three sub-area ROI CTCF measurements per cell type were averaged to yield a representative CTCF value per cell type, per embryo.

Relative spatial fluorescence intensity of tubulin isotypes (presented in Supplementary S3-S8) was assessed in select monolayers (corneal epithelium, lens epithelium and corneal endothelium). Curved cell monolayers were isolated and straightened in ImageJ. Straightened monolayers were oriented along the long axis with basal side down and peripheral to the left, and raw fluorescence intensity matrices (Y x X) were exported as tab-delimited text arrays. In each matrix, the columns (X) represent pixel position along the longitudinal (central/peripheral) axis, and the rows (Y) represent pixel positions along the apicobasal axis. These matrices were utilized for further downstream analysis.

### Statistics and data visualization

All data preparation and statistical analyses were conducted in R (version 4.5.1). Statistical significance was defined as a P-value (or adjusted *P*/FDR where applicable) < 0.05. Significance thresholds were defined as: \**P* < 0.05, \*\**P* < 0.01, \*\*\**P* < 0.001, and ns for *P* ≥ 0.05.

In fluorescence intensity comparisons across cell types, a randomized complete block design was utilized to account for baseline variations in microscopic parameters and fluorescence intensities between independent experimental replicates. Comparisons of tubulin isotype expression across cell types were conducted strictly within stage using a linear mixed effects model (LMM) via the *lme4* package (Bates, Mächler et al. 2015). In this model, “cell type” was designated as the fixed effect while “embryo ID” was designated as the random effect intercept to ensure paired mathematical comparisons within the same biological sample. Estimated marginal means and subsequent pairwise post-hoc comparisons between diverse cell types were performed using the *emmeans* package (Searle 1980). To control for multiple testing across the high dimensional dataset, all raw P-values were adjusted using the Benjamini-Hochberg false discovery rate (FDR) procedure. Graphing was performed using *ggplot2*, *ggh4x*, and *patchwork* packages in R.

For the spatial matrices, matrix coordinates were normalized to unit intervals [0,1]. For apicobasal depth normalization, the Row 1 represented the apical boundary and Row Y_max_ represented the basal lamina. Apicobasal Y score was normalized as:

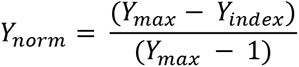

This yielded a score where 1 is apical and 0 is basal.

For the longitudinal axis, two approaches were utilized to compare the early stages (days 3-6) to day 14 as the latter was imaged as a hemi-section. The monolayer length was mapped to an aligned coordinate where for days 3-6:

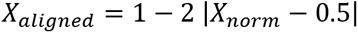

For day 14:

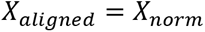

A longitudinal score of 0 denotes a peripheral position and a score of 1 denotes a central position.

Spatial feature extraction was performed to calculate the peak apicobasal modal coordinate (Y_peak_). For the Y_peak_, Y_norm_ was segmented into 100 vertical depth bins. Mean raw intensities were extracted per bin and normalized internally to unit range within each biological replicate to yield relative fluorescence intensity curves. For each biological replicate, the peak apicobasal modal depth (Y_peak_) was identified as the coordinate of maximum intensity:

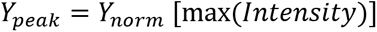

Differences in peak apicobasal position across replicates are presented as the mean modal depth ± standard error of the mean (Y_peak_ ± SEM).

Visualization was done using *ggplot2* and *ggh4x* in R. Replicate-specific profiles were smoothed using local polynomial regression (LOESS) models (span = 0.35-0.45). Within each biological replicate, local fit uncertainty was visualized through standard error ribbons surrounding smoothed curves.

## ACKNOWLEDGEMENTS

The authors acknowledge the members of the Rogers Laboratory and the Comparative Ophthalmology and Vision Sciences Laboratory at UC Davis School of Veterinary Medicine for their contributions throughout the study period. Special appreciation to the UC Davis Hopkins Avian Facility for their continued support.

## AUTHOR CONTRIBUTIONS

Conceptualization: R.R, W.R.S., C.D.R., S.M.T.; Data curation: R.R, W.R.S., M.M., S.B..; Formal analysis: R.R, W.R.S.; Funding acquisition: R.R., C.D.R., S.M.T.; Methodology: R.R, W.R.S.; Project administration: R.R., C.D.R., S.M.T.; Resources: C.D.R., S.M.T.; Supervision: R.R, W.R.S., C.D.R., S.M.T.; Visualization: R.R,

W.R.S., M.M.; Writing – original draft: R.R, W.R.S. C.D.R.; Writing – review & editing: R.R, W.R.S., C.D.R., S.M.T.

