## Supplemental Figures for "A developmental resource mapping tubulin isotype deployment during corneal morphogenesis"

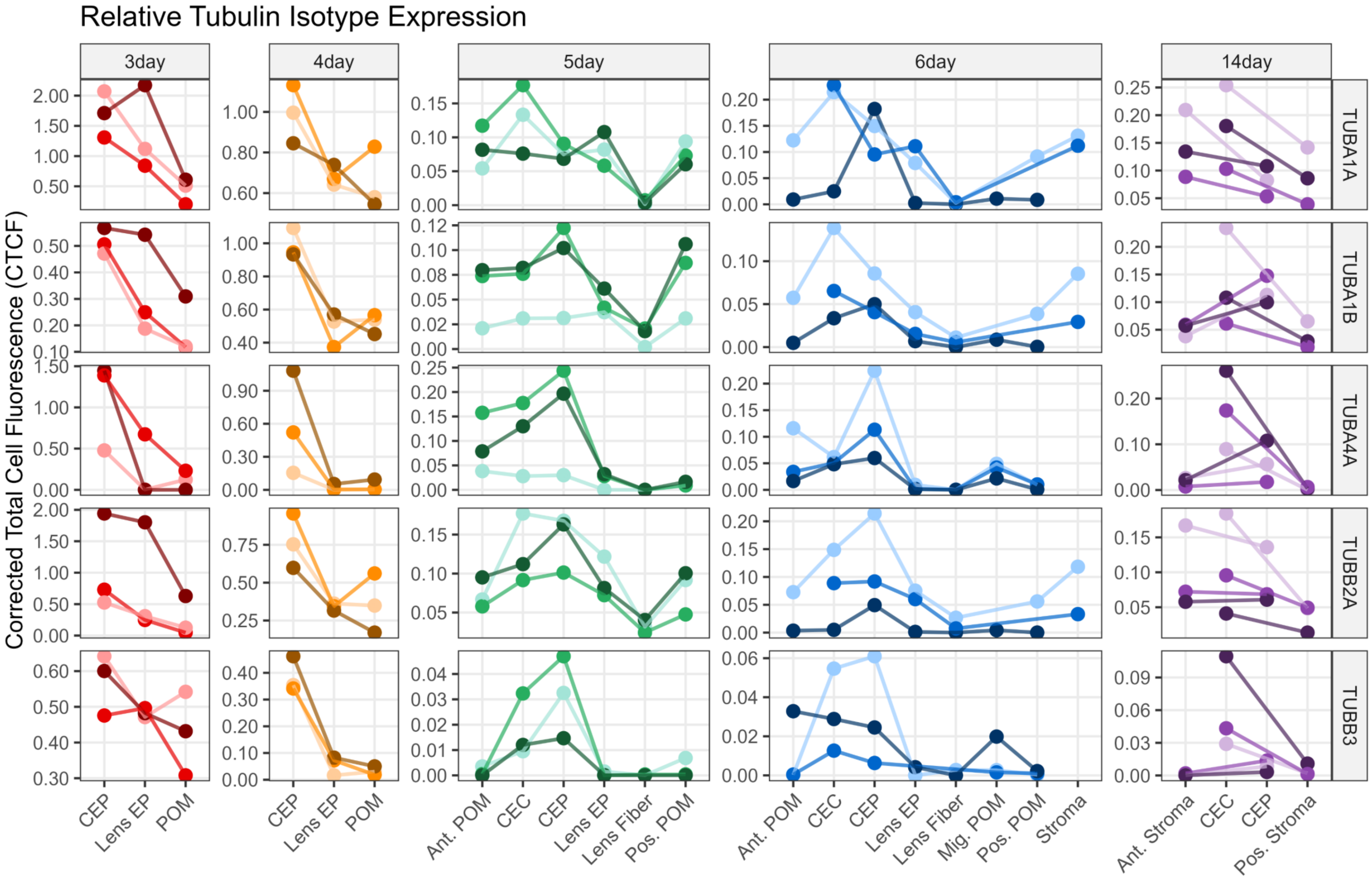


**Fig. S1. Global statistical summary of tubulin isotype differential expression across developing chick anterior ocular tissues.** Composite heatmap displaying pairwise statistical significance of cell type comparisons across all tubulins isotypes investigated across all stages. Pairwise differences were modelled using a linear mixed effects model. Color gradient reflects -$\log_{10}$ transformed false discovery rate (FDR) adjusted P-values. Darker red indicates highly significant differences in cell type tubulin isotype fluorescence intensity expressed as corrected total cell fluorescence. Asterisks indicate significance thresholds (* FDR < 0.05; ** FDR < 0.01; *** FDR < 0.001). ns, not significant (FDR >= 0.05).

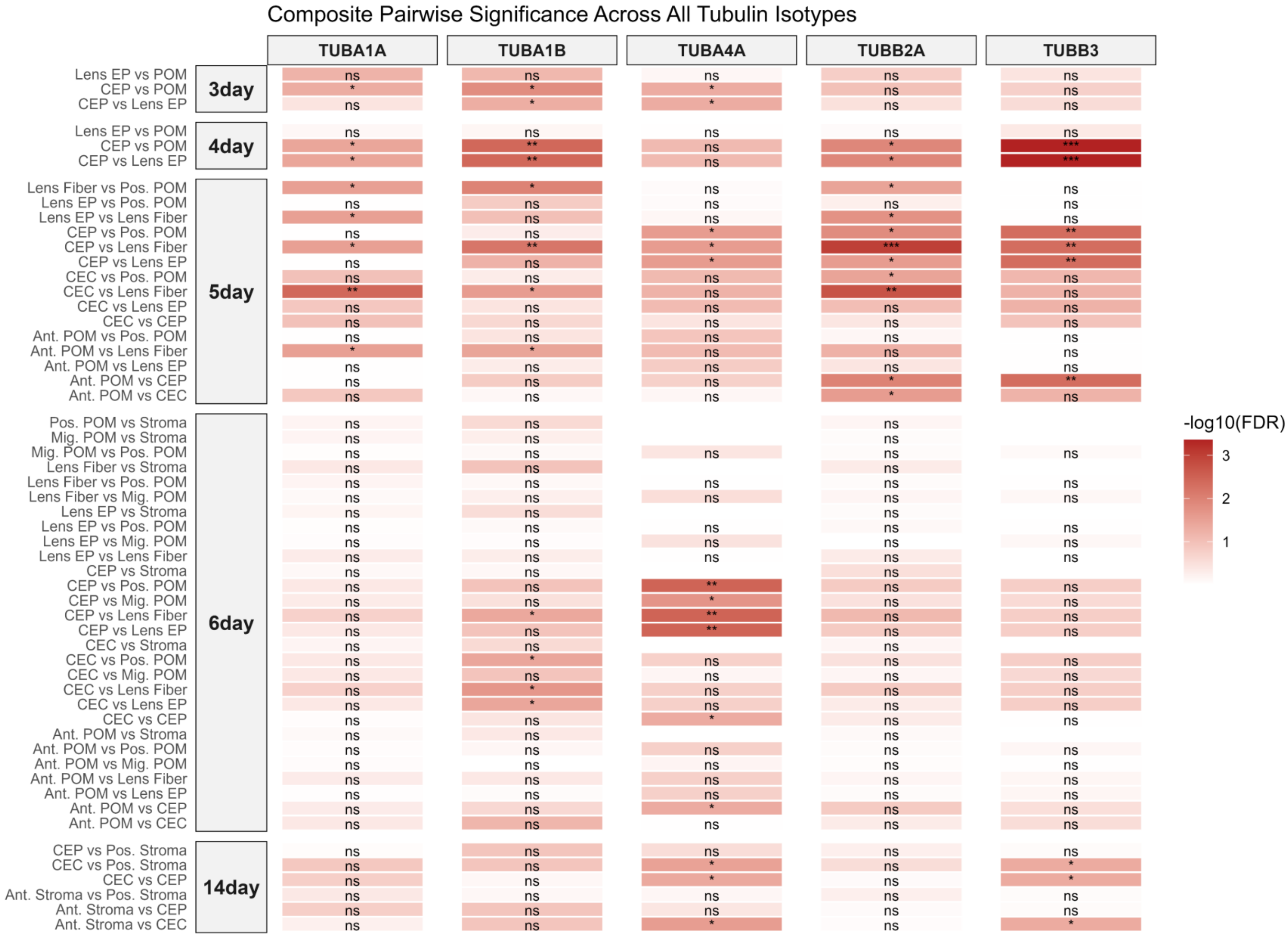


**Fig. S2 Developmental expression dynamics of select tubulin isotypes across developing chick anterior ocular tissues.** Relative fluorescence intensity of tubulin isotypes quantified via ImageJ across developmental stages and represented by corrected total cell fluorescence (CTCF). Individual data points represent the mean CTCF of three subarea measurements per cell type within a single biological replicate (N = 3). Connecting lines denote paired observations from the same biological replicate. Y axes are scaled independently to illustrate cell type CTCF variation. CEP, corneal epithelium; CEC, corneal endothelium; Lens EP, lens epithelium; POM, periocular mesenchyme; Ant., anterior; Pos., posterior; Mig., migratory.

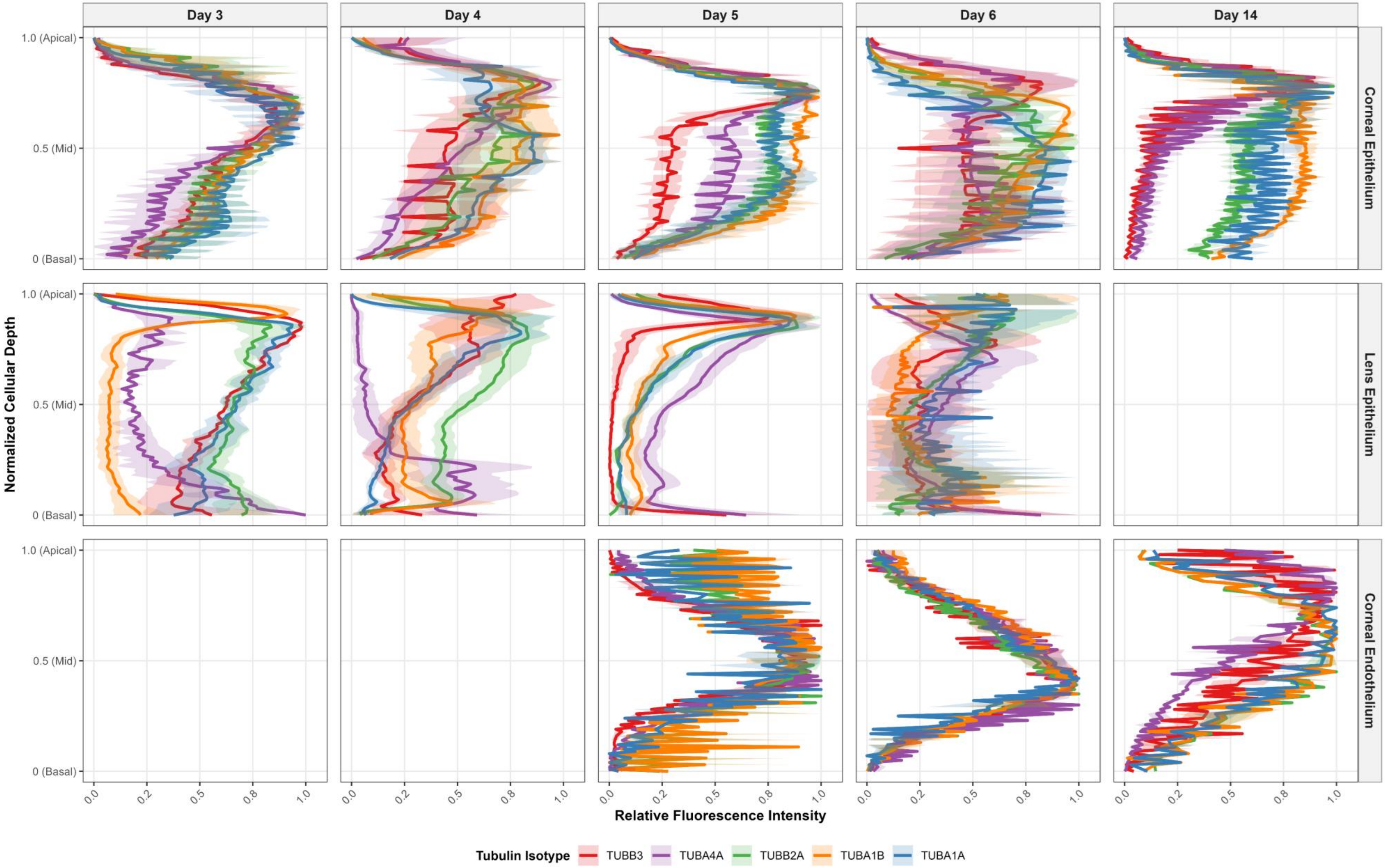


**Fig. S3. Apicobasal fluorescence localization profiles of all tubulin isotypes assessed in study across chick embryonic ocular monolayers.** Relative fluorescence intensity of all tubulin isotypes investigated, quantified via ImageJ within straightened monolayers across developmental stages along the normalized apicobasal cell axis. Colored curves represent local regression (LOESS) fits for average distribution of each isotype (N = 3), flanked by shaded standard error ribbons indicating fit uncertainty within each isotype across samples.

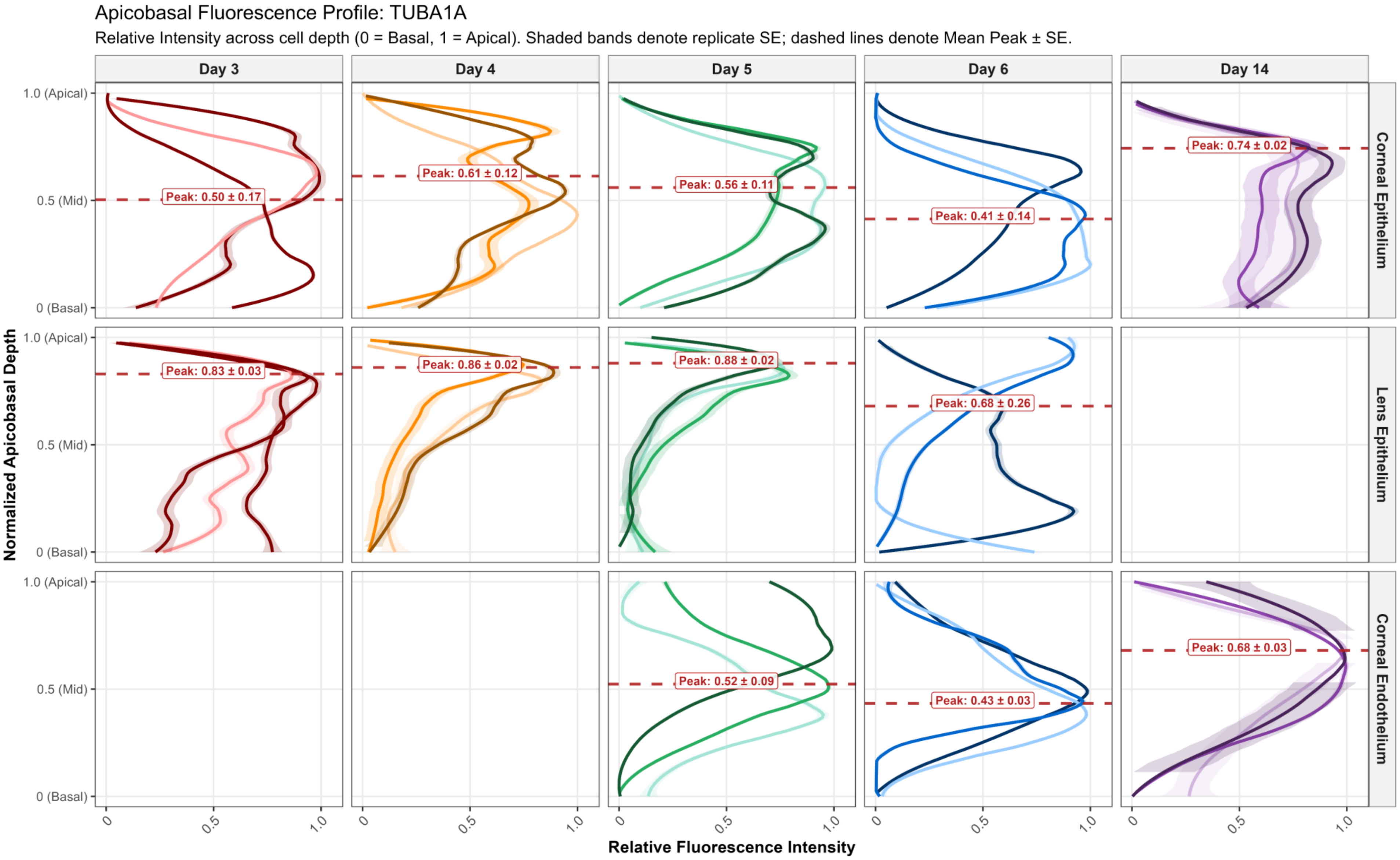


**Fig. S4. Apicobasal fluorescence localization profiles of TUBA1A across chick embryonic ocular monolayers.** Relative fluorescence intensity of TUBA1A quantified via ImageJ within straightened monolayers across developmental stages along the normalized apicobasal cell axis. Colored curves represent local regression (LOESS) fits for individual biological replicates (N = 3), flanked by shaded standard error ribbons indicating fit uncertainty within each sample. Red horizontal dashed lines denote mean peak apicobasal depth ($Y_{peak}$) with numeric values annotated (mean ± SEM).

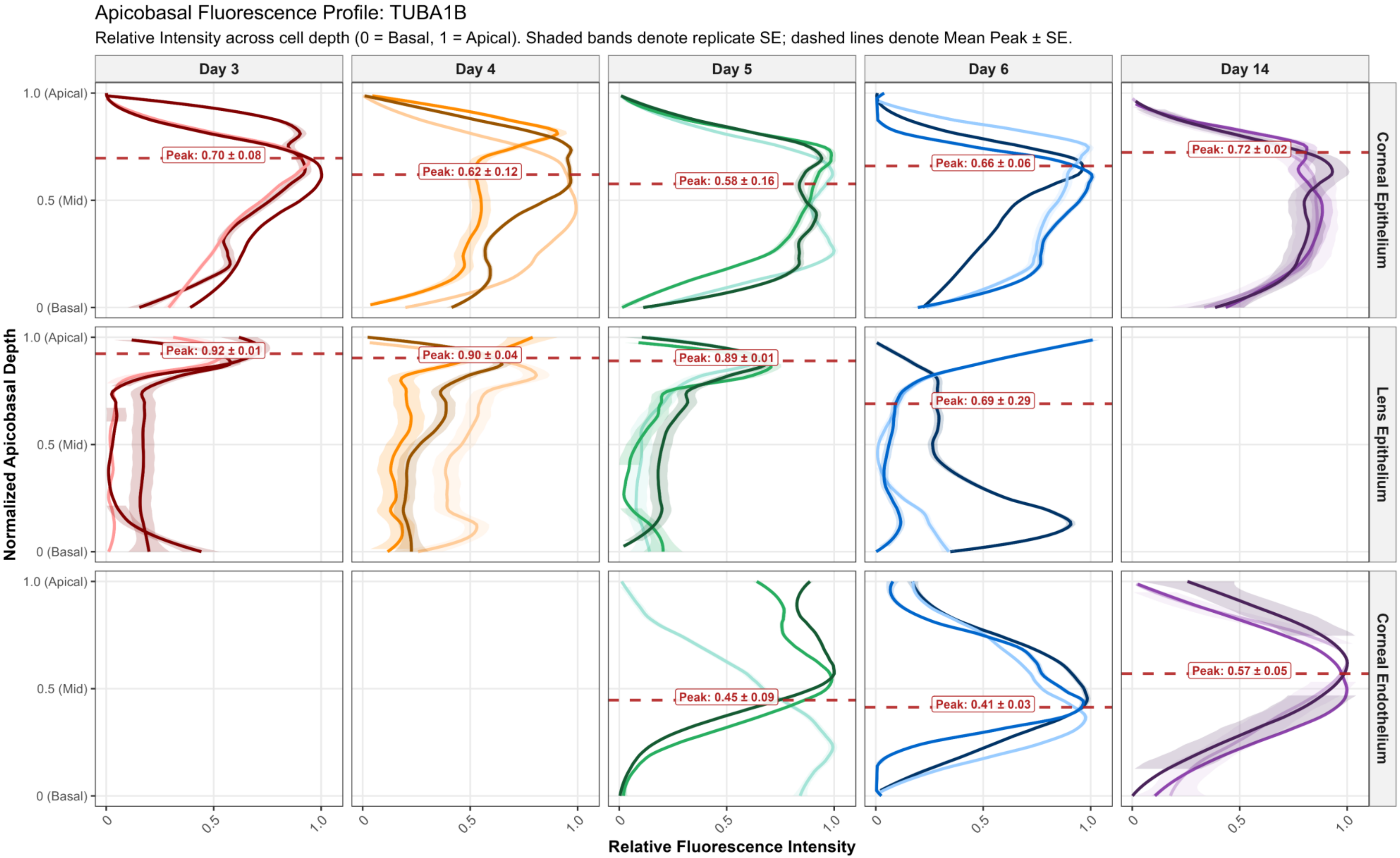


**Fig. S5. Apicobasal fluorescence localization profiles of TUBA1B across chick embryonic ocular monolayers.** Relative fluorescence intensity of TUBA1B quantified via ImageJ within straightened monolayers across developmental stages along the normalized apicobasal cell axis. Colored curves represent local regression (LOESS) fits for individual biological replicates (N = 3), flanked by shaded standard error ribbons indicating fit uncertainty within each sample. Red horizontal dashed lines denote mean peak apicobasal depth ($Y_{peak}$) with numeric values annotated (mean ± SEM).

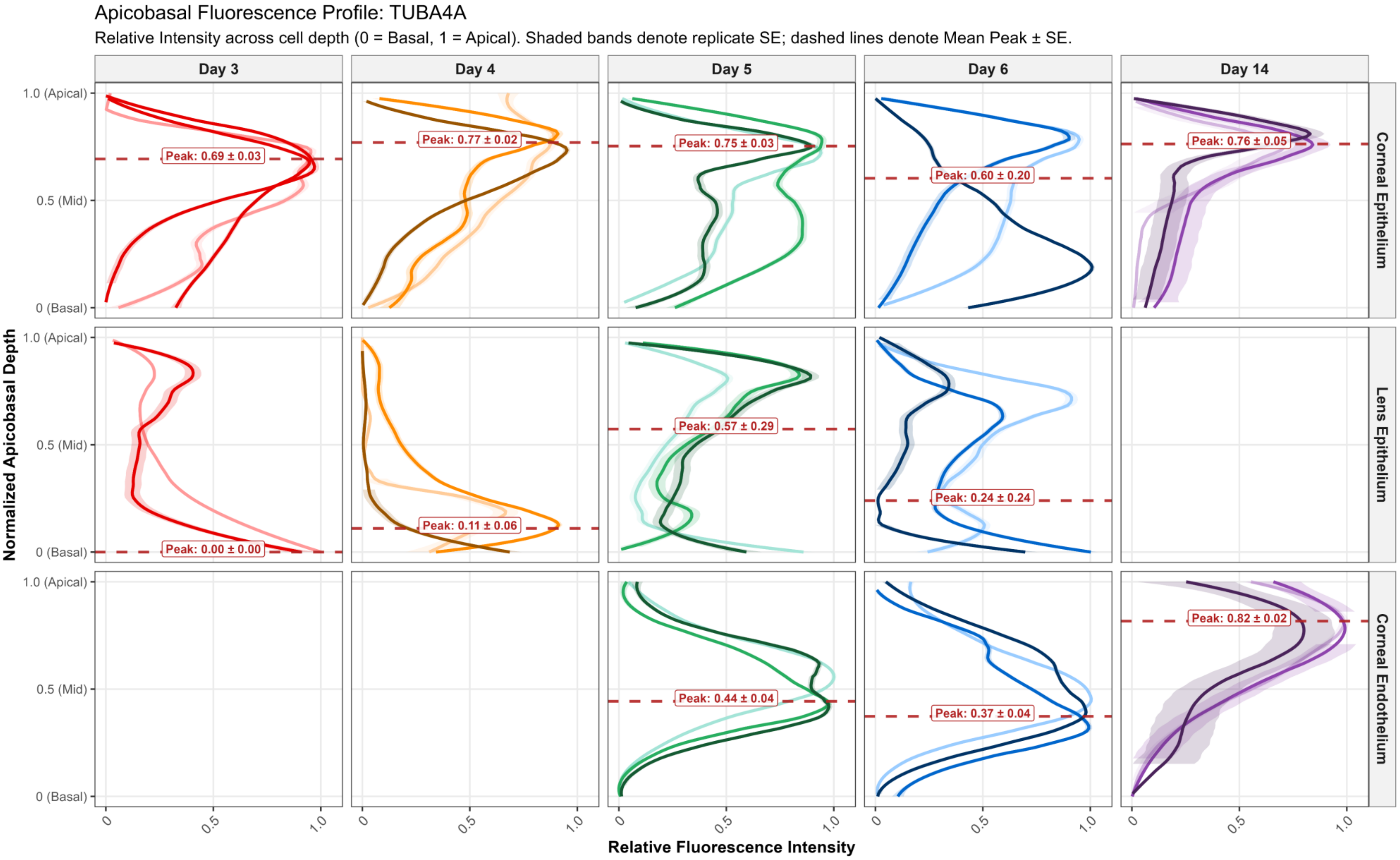


**Fig. S6. Apicobasal fluorescence localization profiles of TUBA4A across chick embryonic ocular monolayers.** Relative fluorescence intensity of TUBA4A quantified via ImageJ within straightened monolayers across developmental stages along the normalized apicobasal cell axis. Colored curves represent local regression (LOESS) fits for individual biological replicates (N = 3), flanked by shaded standard error ribbons indicating fit uncertainty within each sample. Red horizontal dashed lines denote mean peak apicobasal depth ($Y_{peak}$) with numeric values annotated (mean ± SEM).

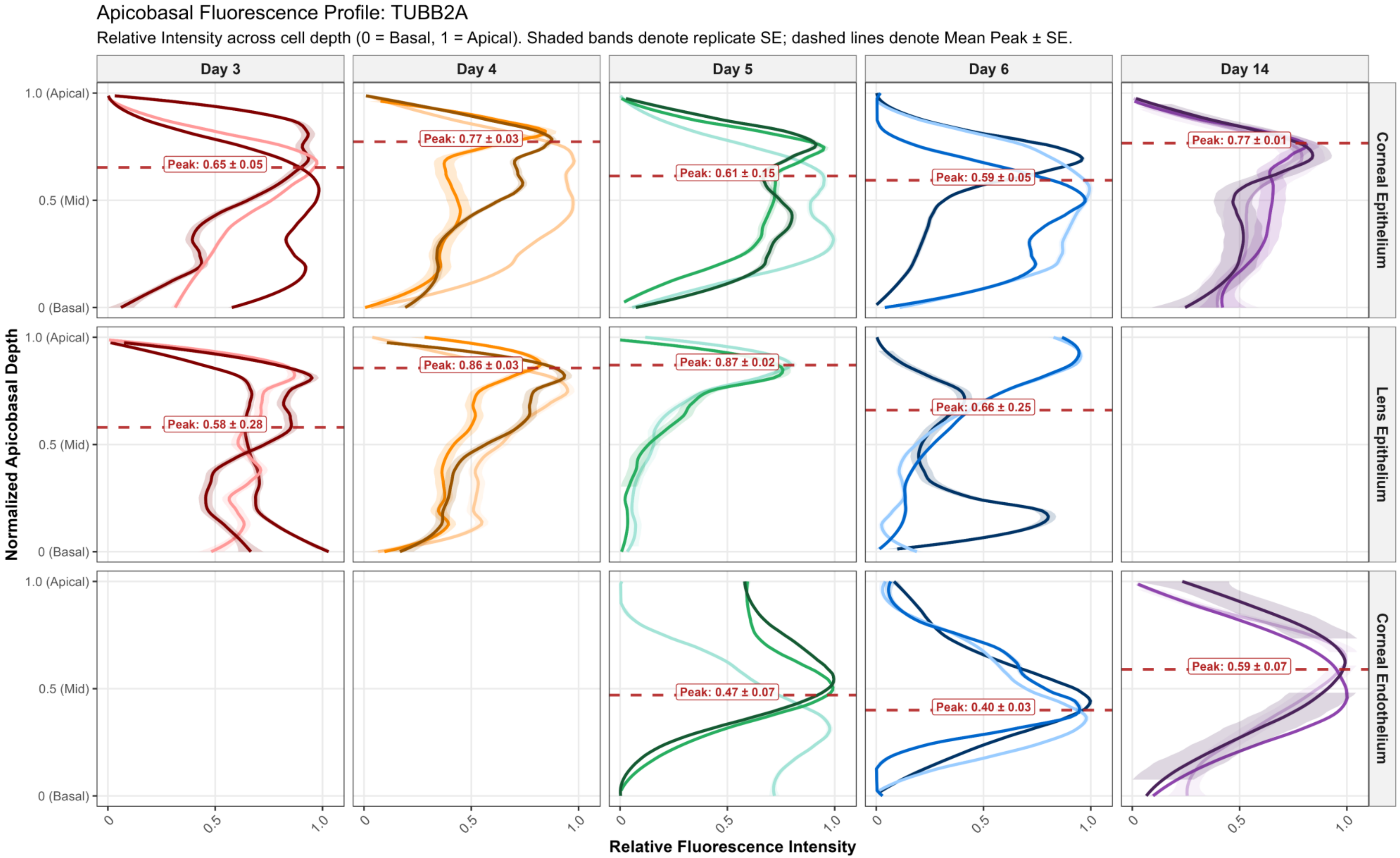


**Fig. S7. Apicobasal fluorescence localization profiles of TUBB2A across chick embryonic ocular monolayers.** Relative fluorescence intensity of TUBB2A quantified via ImageJ within straightened monolayers across developmental stages along the normalized apicobasal cell axis. Colored curves represent local regression (LOESS) fits for individual biological replicates (N = 3), flanked by shaded standard error ribbons indicating fit uncertainty within each sample. Red horizontal dashed lines denote mean peak apicobasal depth ($Y_{peak}$) with numeric values annotated (mean ± SEM).

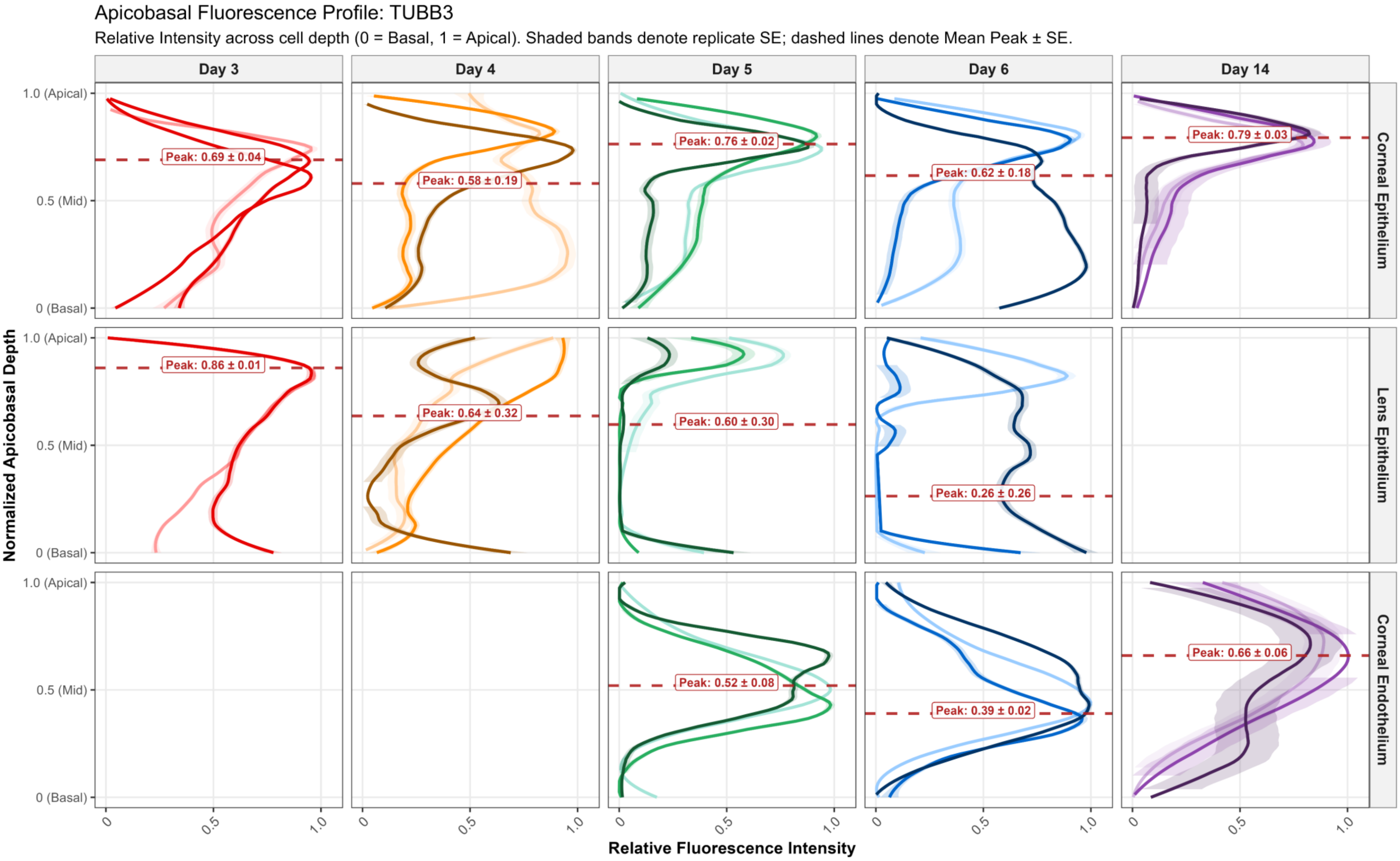


**Fig. S8. Apicobasal fluorescence localization profiles of TUBB3 across chick embryonic ocular monolayers.** Relative fluorescence intensity of TUBB3 quantified via ImageJ within straightened monolayers across developmental stages along the normalized apicobasal cell axis. Colored curves represent local regression (LOESS) fits for individual biological replicates (N = 3), flanked by shaded standard error ribbons indicating fit uncertainty within each sample. Red horizontal dashed lines denote mean peak apicobasal depth ($Y_{peak}$) with numeric values annotated (mean ± SEM).
